# Historical dispersal and host-switching formed the evolutionary history of a globally distributed multi-host parasite - the *Ligula intestinalis* species complex

**DOI:** 10.1101/2022.10.12.511855

**Authors:** Masoud Nazarizadeh, Milena Nováková, Géraldine Loot, Nestory P. Gabagambi, Faezeh Fatemizadeh, Odipo Osano, Bronwen Presswell, Robert Poulin, Zoltán Vitál, Tomáš Scholz, Ali Halajian, Emiliano Trucchi, Pavlína Kočová, Jan Štefka

**Affiliations:** Faculty of Science, University of South Bohemia, České Budějovice, Czech Republic; Institute of Parasitology, Biology Centre CAS, České Budějovice, Czech Republic; UMR-5174, EDB (Laboratoire Evolution and Diversité Biologique), CNRS, IRD, Université Toulouse III Paul Sabatier, France; Tanzania Fisheries Research Institute, Kyela, Mbeya, Tanzania; Department of Environmental Science, Faculty of Natural Resources, University of Tehran, Karaj, Iran; School of Environmental Studies, University of Eldoret, Kenya; Department of Zoology, University of Otago, Dunedin, New Zealand; Research Center for Fisheries and Aquaculture, Institute of Aquaculture and Environmental Safety, Hungarian University of Agriculture and Life Sciences, Szarvas, Hungary; Research Administration and Development, and 2-DSI-NRF SARChI Chair (Ecosystem health), Department of Biodiversity, University of Limpopo, South Africa; Department of Life and Environmental Sciences, Marche Polytechnic University, Ancona, Italy; Čakovice, Týnec nad Sázavou, Czech Republic

**Keywords:** Historical biogeography, demographic history, *Ligula* species complex, host specificity

## Abstract

Studies on parasite biogeography and host spectrum provide insights into the processes driving parasite diversification. Global geographical distribution and a multi-host spectrum make the tapeworm *Ligula intestinalis* a promising model for studying both the vicariant and ecological modes of speciation in parasites. To understand the relative importance of host association and biogeography in the evolutionary history of this tapeworm, we analysed mtDNA and reduced-represented genomic SNP data for a total of 139 specimens collected from 18 fish-host genera across a distribution range representing 21 countries. Our results strongly supported the existence of at least 10 evolutionary lineages and estimated the deepest divergence at approximately 4.99-5.05 Mya, which is much younger than the diversification of the fish host genera and orders. Historical biogeography analyses revealed that the ancestor of the parasite diversified following multiple vicariance events and was widespread throughout the Palearctic, Afrotropical, and Nearctic between the late Miocene and early Pliocene. Cyprinoids were inferred as the ancestral hosts for the parasite. Later, from the late Pliocene to Pleistocene, new lineages emerged following a series of biogeographic dispersal and host-switching events. Although only a few of the current *Ligula* lineages show narrow host-specificity (to a single host genus), almost no host genera, even those that live in sympatry, overlapped between different *Ligula* lineages. Our analyses uncovered the impact of historical distribution shifts on host switching and the evolution of host specificity without parallel host-parasite co-speciation. Historical biogeography reconstructions also found that the parasite colonized several areas (Afrotropical and Australasian) much earlier than was suggested by only recent faunistic data.

## Introduction

Host specificity is a key feature of parasitic and symbiotic organisms. It defines their ability to survive on different hosts (Poulin, 2011), disperse (Mácová et al., 2018) and maintain the genetic diversity of populations (Martinu et al., 2018; Wacker et al., 2019). The evolution of host specificity in multi-host parasites is a prospective candidate for the mechanism of sympatric adaptive speciation (Kalbe et al., 2016). For long, the evolution of host-specific lineages has been viewed mostly from the co-speciation (or co-diversification) angle, where the evolution of the parasite is directly associated with that of the host(s) (Hafner & Nadler, 2017; Štefka et al., 2011). However, the origin of host-specific lineages is often linked to a series of historical biogeographical events (Ding et al., 2022; Perrot-Minnot et al., 2018). Therefore, incongruence may appear between the parasite and host phylogenies for various reasons, including founder events, bottlenecks and extinctions either in the host or parasite, and, most importantly, due to host-switching that could have occurred over large-scale episodic periods of palaeo-environmental and climatic fluctuations, such as Quaternary glaciations (Hoberg et al., 2012; Johnson et al., 2002; Zarlenga et al., 2006). Under stable environmental conditions, co-divergence between closely associated organisms may occur (Haukisalmi et al., 2016), whereas under changing conditions, hosts possibly move in search of suitable habitats and find themselves in contact with species they did not previously overlap with. These fluctuations provide new opportunities for parasites to switch between host species (Hoberg et al., 2012; Hoberg & Brooks, 2008, 2013).

Following a successful host switch, gene flow may be maintained or ceased between the ancestral and newly established parasite population, depending on the duration of the distribution overlap between the host species (e. g., Techer et al., 2022; Wang et al., 2016). Phylogenomic and historical biogeography methods allow for reconstructing past distribution changes and associated gene flow events in parasite lineages, providing tools to study the emergence and diversification of parasite species, identification of cryptic lineages and prediction of disease transmission (Angst et al., 2022; Doyle et al., 2022). Yet, although metazoan parasites represent 40% of all living metazoan species, relatively few studies have investigated the historical biogeography of parasitic species (Dobson et al., 2008). Instead, these studies often investigate the host–parasite co-phylogeny. Whilst the co-phylogenetic approach has demonstrated significant co-evolutionary signals in parasites with a direct life cycle (Rahmouni et al., 2022; Šimková et al., 2004, 2022), it often finds weak host–parasite co-evolutionary signals in generalist multi-host parasites or parasites with complex life cycles involving intermediate hosts, possessing more varied dispersal opportunities (Hoberg & Brooks, 2008; Perrot-Minnot et al., 2018). Importantly, the co-phylogenetic framework does not consider the evolution of host-specific lineages in parasites whose origin is much younger than the age of the hosts, where the co-phylogenetic methods simply cannot be applied (Bouzid et al., 2008).

To study the contribution of past processes to the evolution of host-specific parasite lineages, we applied historical biogeography methods to study the tapeworm *Ligula intestinalis* sensu lato (Cestoda: Diphyllobothriidea). Its widespread geographical distribution and multi-host spectrum make it a promising model for studying the vicariant and ecological modes of speciation (Hoole et al., 2010; Nazarizadeh et al., 2022a; Štefka et al., 2009). It is an obligatory endoparasite with a complex life cycle involving two intermediate hosts (crustacean copepod as the first and planktivorous fish as the second intermediate host) and birds feeding on the infected fish as the definitive host (Dubinina, 1980). *L. intestinalis* spends a maximum of only five days in its final host, rapidly entering the sexual phase and dispersing eggs into the water with host faeces, while the secondary larval stage, plerocercoid, represents the most distinct phase of the parasite’s life cycle (Dubinina, 1980). It lasts at least 10 months and occurs in the peritoneal cavity of planktivorous fish, negatively impacting the health, fertility and behaviour of individual fish (Gabagambi et al., 2019; Loot et al., 2002) as well as the ecology of the entire fish population (Kennedy et al., 2001). *L. intestinalis* usually appears in cyprinoid fish [previously the family Cyprinidae], but is capable of invading intermediate hosts from various taxonomic groups such as Salmonidae, Galaxiidae and Catostomidae (Chapman et al., 2006; Dubinina, 1980; Štefka et al., 2009).

Studies exploring the phylogenetic relationships and population structure of the *L. intestinalis* species complex identified at least six major lineages with different levels of host and geographic distribution: (1) clade A (European and north African populations), (2) clade B (European, Chinese, Australian and North African populations), (3) *L. alternans* (Asia) [previously *L. digramma*], (4) Canada, (5) Chinese and (6) Ethiopian populations (Bouzid et al., 2008; J. Štefka et al., 2009). The studies indicated that both geography and host specificity may determine the genealogical relationships of the parasites by inducing genetic differentiation. However, their sampling lacked several biogeographical areas, and the historical demography of the lineages was not elucidated. Thus, the course of interaction between the historical geographical distribution of the parasite and the evolution of host-specific lineages has remained a contentious issue.

The main objective of the present study is to reveal the contribution of historical geographical distribution and past host usage (specificity) in the formation of contemporary lineage diversity in parasites with high dispersal capabilities. Using a geographically wide sampling of *Ligula* tapeworms as a study system, we characterised it with SNP genotyping and mitochondrial sequencing. We aimed to i) determine the true lineage diversity and levels of host specificity of parasites forming a species complex which lacks distinctive morphological traits; ii) provide dates for major diversification events in the evolutionary history of the parasite, and iii) determine the factors and events influencing lineage diversity in space and time by the analyses of historical areas, demographic history of lineages and past gene flow.

## Materials and Methods

### Sample collection and DNA extraction

We analysed 139 specimens of *Ligula* plerocercoids collected from 18 fish genera over a broad geographic area representing 21 countries (Table S1). Integrity and quantity of newly isolated samples (extracted with DNeasy blood and tissue kit [Qiagen] from specimens preserved in 96% ethanol) and DNA extracts from previous studies (Bouzid et al., 2008; Štefka et al., 2009) were verified on 0.8% agarose gel and with Qubit 2.0 Fluorometer (Invitrogen). The final set of mitochondrial (86 new specimens were added to 76 sequences from Bouzid et al., 2008 and Štefka et al., 2013 studies) and ddRAD (all new specimens) samples covered the distribution of the major lineages found in previous studies, which was complemented with several new areas to represent five terrestrial biogeographic realms: Nearctic, Palearctic, Afrotropical, Oriental and Australasian (Fig. 1, Table S1a and S1b).

**Fig. 1.**
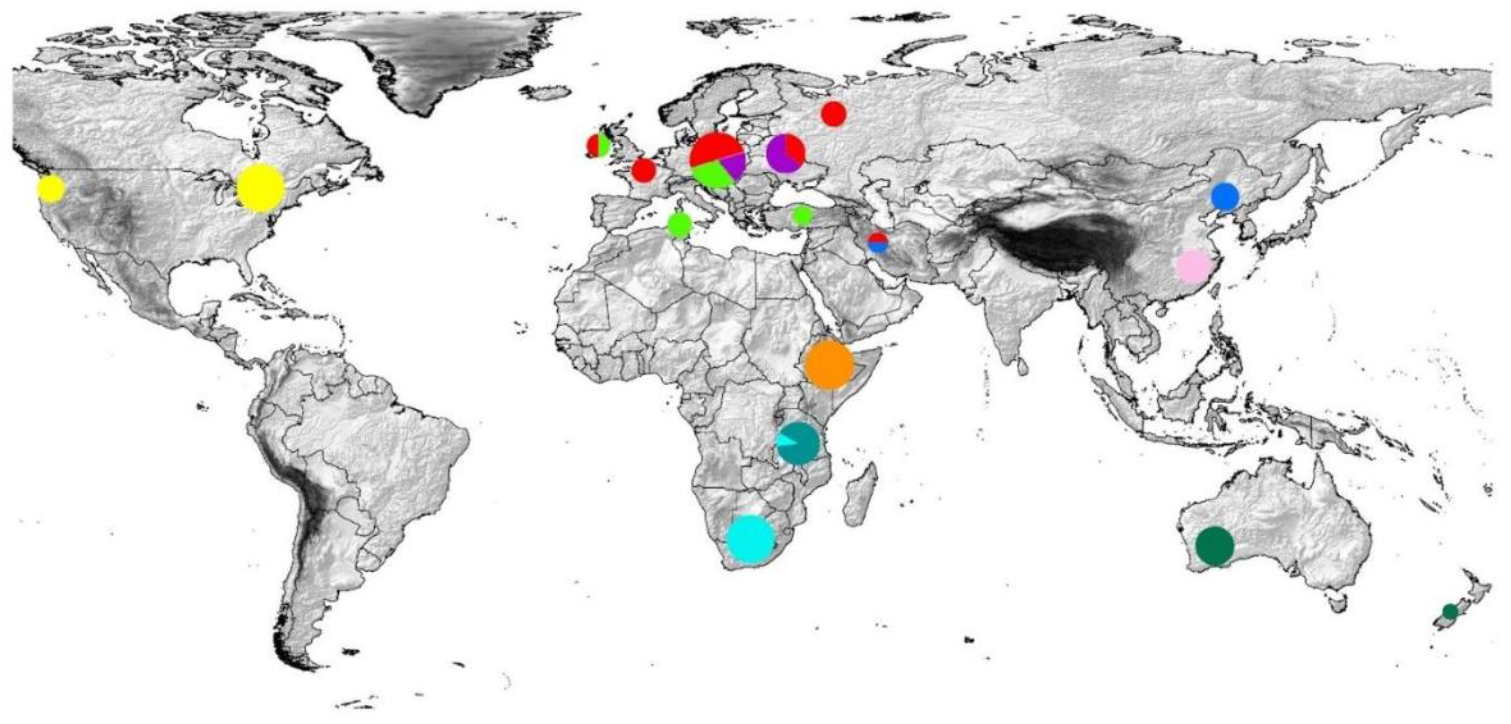
*Ligula* sampling localities and patterns of genetic structure. Pie charts reflect the proportion of individuals assigned to each phylogenetic lineage (shown in Fig. 2); Lineage A (green), Lineage B (red), *L. alternans* (blue), China (light pink), Australia and New Zealand (dark green), *L. pavlovskii* (magenta), East African Rift (EAR; dark turquoise), Central and South Africa (CSA; light turquoise), Ethiopia (orange), and Nearctic (yellow).

### Mitochondrial DNA sequencing

Amplification of three fragments of the mitochondrial genome, including cytochrome b (Cyt *b*), cytochrome oxidase subunit 1 (COI) and NADH dehydrogenase 1 (ND1) genes generated a concatenated dataset with a total length of 1693 bp. All genes were amplified by polymerase chain reaction (PCR; see Supplementary Material, section 1) and sequenced using primers specifically designed for this study or previously published primers (Table S2). Sanger sequences were assembled and edited using Geneious Prime v.2022.1 (Gene Codes) and checked for possible stop codons based on the Alternative Flatworm Mitochondrial Code (transl_table 14).

### ddRAD library preparation and reference genome

ddRAD libraries were prepared using 350 ng of DNA per sample following a modified version of the protocol proposed by Peterson et al. (2012) (Supplementary Material, section 2). Two samples of *Dibothriocephalus latus* (Diphyllobothriidea) were included as outgroups. Libraries were sequenced at several lanes of Illumina NovaSeq 150 bp PE (Novogene UK) yielding 6.8 million paired-end reads per sample on average. Steps for preparation of multiplexed ddRAD-seq libraries for each barcoded individual sample (Table S1a) and purification are outlined in supplementary material, section 2. We also generated a draft genome of *Ligula* using Illumina platforms (Omniseq), yielding an assembly comprising 49,218 scaffolds and covering 780 Mb of the genome; gene prediction and annotation for this assembly are ongoing.

### ddRAD data assembly and SNP calling

FastQC (Andrews, 2010) was applied for an initial quality check. The process_radtags program in Stacks v.2.5.3 was used to demultiplex and filter the raw reads with low quality and uncalled bases (Rochette et al., 2019). Also, cut sites, barcodes and adaptors were removed from ddRAD outputs. Next, paired-end read alignment of each sample to the reference genome was performed using the default mode of Bowtie 2 (Langmead & Salzberg, 2012). SNPs calling and assembling loci were carried out from the alignment using the ref_map.pl wrapper program. Data were then filtered for missingness, minor allele frequency (MAF) and linkage disequilibrium (LD) in Stacks (see Supplementary Material, section 3 for details). In sum, we generated three SNP matrices in variant call format (VCF) comprising (1) 44,298 SNPs with a mean coverage per locus of ca. 20×, (2) second linkage disequilibrium filtered data set (0.2 squared coefficient of correlation) including 10,937 SNPs and a mean coverage per locus of ca. 18×, and (3) two outgroups which were added to the third data set comprising 4471 unlinked SNPs and a mean coverage per locus of ca. 25×.

### Analysis of sequence diversity

Three gene fragments (Cyt *b*: 405 bp COI: 396 bp, and ND1: 893 bp) were added to an alignment that included 76 sequences of Cyt *b* and COI from Bouzid et al. (2008), Bouzid et al. (2013) and Bouzid et al. (2013) studies. All gene fragments were aligned separately using MUSCLE performed in MEGA v.5 (Tamura et al., 2011). We used the three gene fragments for phylogenetic analyses; however, all genetic distance-based analyses were carried out based on Cyt *b* and COI sequences which had no missingness. Analysis of population genetic diversity (sequence polymorphism, haplotype diversity and nucleotide diversity) was performed in DnaSP v.5 (Librado & Rozas, 2009) on samples grouped according to their phylogenetic relationships (see below).

For nuclear SNP data, we estimated the expected (HE) and observed (HO) heterozygosities, private alleles, and inbreeding coefficients among the parasite populations using the R package SambaR (de Jong et al., 2021). The first ddRAD dataset (44,298 SNPs) was applied for the analysis as the calculation assumed that the input data were not reduced based on LD filtering. We used Plink v.1.90 (Chang et al., 2015) to convert the VCF file to the bed format. The default data filtering was applied with the “filterdata” function in the R package (see De Jong et al., 2021).

### Phylogenetic analyses

The best-fitting substitution models and partition schemes were estimated for the three mtDNA genes using a greedy algorithm in PartitionFinder v.2.1.1 (Lanfear et al., 2017). Using Bayesian inference (BI) and maximum likelihood (ML), the phylogenetic relationships of the *Ligula* lineages were reconstructed. BI analysis was conducted using MrBayes v.3.1.2 (Ronquist & Huelsenbeck, 2003) on the data partitioned based on codon positions and using the settings as described in supplementary material, section 4. We inferred the ML tree using IQTREE v.2.1.2 (Minh et al., 2020) under the best-fitting model from PartitionFinder (Table S3) and 1,000 ultrafast bootstrap replicates. We also used the -bnni option to minimize the risks of overestimating support values to a minimum. We included *Dibothriocephalus nihonkaiensis* and *D. latus* (Li et al., 2018a), sister groups of *Ligula*, as the outgroups.

We utilised the third data set (4471 unlinked SNPs) to reconstruct the phylogenetic gene tree and the species tree using three approaches. Using PartitionFinder, we first determined the parameters applied as input into RAxML-ng to reconstruct an ML tree (Kozlov et al., 2019). Second, the alignment was used for a Bayesian phylogenetic analysis. BI was completed using ExaBayes v.1.5 (Aberer et al., 2014) under a model partitioning scheme analogous to the ML analyses. Third, species trees were investigated using SVDquartets in PAUP v.4.0a147 (Chifman & Kubatko, 2014). Further details on the assessments of convergence and ML, BI and species trees are provided in supplementary material, section 4.

### Population structure analyses

Using the concatenated COI and Cyt *b* (801 bp), haplotype networks were constructed to visualize relationships among the parasite populations via the Median-Joining algorithm in PopArt v.1.7 (Leigh & Bryant, 2015). Moreover, ddRAD data set was applied to construct a phylogenetic network using a neighbour-net algorithm and 1,000 bootstraps executed in SplitsTree v.4.10 (Huson & Bryant, 2006).

Several clustering analyses were performed to discover the number of distinct genetic groups. First, the model-based evolutionary clustering method was applied in ADMIXTURE (Alexander et al., 2009) via its parallel processing capabilities (AdmixPiPe: Mussmann et al., 2020). Second, a Principal Component Analysis (PCA) was performed using the glPCA function from the adegenet R package (Jombart & Collins, 2015). Next, we applied the TESS3R package in R to provide additional estimates of the role of geography in the genetic structure (Caye et al., 2018). TESS3R explores the implications of genetic diversity in natural populations using geographic and genotypic data simultaneously. Lastly, we applied the programs fineRADstructure and RADpianter v.0.2 (Malinsky et al., 2018) to reveal population genetic structure based on the nearest-neighbour haplotype (For details on ADMIXTURE and fineRADstructure analyses, see Supplementary Material, section 5).

### Species delimitation analyses

To detect species boundaries in the *Ligula* species complex using the mtDNA data, species delimitation was performed using three independent approaches (genetic distance and tree-based approaches): the Generalized Mixed Yule Coalescent model (GMYC by Pons et al., 2006), Bayesian implementation of Poisson Tree Processes model (bPTP by Zhang et al., 2013), and Assemble Species by Automatic Partitioning (ASAP by Puillandre et al., 2021 Supplementary Material, section 6).

To make comparisons among species delimitation models using SNP data, we conducted Bayes Factor Delimitation (BFD, Leaché et al., 2014) implemented in SNAPP in BEAST2 (Bouckaert et al., 2014; Bryant et al., 2012). Comparing models by marginal likelihood scores, this approach assesses support for the alternative model using Bayes factors under the multispecies coalescent (MSC) model. We tested a nested set of hypotheses up to the maximum potential number of species, where every lineage was considered a different species (Supplementary Material, section 6).

### Divergence time and phylogeographic structure

For mtDNA, divergence dates were estimated using a coalescent-based model implemented in BEAST v.1.8.2 (Drummond et al., 2012) Due to a lack of available fossil or geographic events, we used two common rates of substitution for the mt-genes (Cyt *b* =0.0195, COI= 0.0225 substitutions/site/Myr), which were previously estimated for species of *Taenia* tapeworms (Cyclophyllidea). In addition, we applied the divergence time between *Spirometra* and *Diphyllobothrium* genera (at the base of Diphyllobothriidea), which is normally distributed with a mean of 11.47 Mya, a standard deviation of 1.14 Mya and a 95% interval of 9.58–13.36 Mya. The best-fit partitioning schemes were assigned based on the Akaike information criterion (AICc) in PartitionFinder and the analysis was run using the settings reported in the supplementary material, section 7.

To infer the divergence time among major tapeworm lineages using SNP data, we first estimated the substitution rates for protein-coding sequences (CDS) of the *L. intestinalis* genome based on the methods suggested by Wang et al. (2015). Protein sequences of 10 flatworms (*D. latus, Spirometra erinaceieuropaei, Taenia solium, T. asiatica, T. saginata, Echinococcus granulosus, E. multilocularis, Hymenolepis microstoma, Schistosoma japonicum, S. mansoni*) were obtained from WormBase (https://wormbase.org/). After concatenating all CDS parts, they were aligned together using MAFFT v.7.4 (‘L-INS-I’ algorithm) (Katoh & Standley, 2013) and their gaps were discarded by Trimal v.1.4 (Capella-Gutiérrez et al., 2009). The best evolution models for the CDS alignments were calculated using PartitionFinder (GTR+I+G). ML trees were reconstructed with 400 bootstrapping replicates using RAxML under the best fit models. The substitution rate (branch-length) for *Ligula* was estimated to be 0.0201 which was 1.02-fold higher than that of *T. solium* (0.0196 mutations per site) (Fig. S1). Wang et al. (2015) calculated the mutation rate/site/year in the *T. solium* genome (2.82 × 10^−9^). Therefore, we estimated 2.89 × 10^−9^ substitutions/site/year for the *Ligula* genome. We used Beast v1.8 with 141 sequences, including two *D. latus* outgroups to reconstruct the divergence dates of the sequences under a birth-death process as a tree prior and the uncorrelated relaxed molecular clock model for the mutation rate (2.89 × 10^−9^). The analysis was run using the settings presented in the supplementary material, section 7.

### Historical biogeography and ancestral host reconstructions

Inferring ancestral ranges and the historical biogeography of *Ligula* was done based on three models of biogeographical range extension, including Bayesian inference (BAYAREA-like), Dispersal-Vicariance (DIVA-like), and Dispersal-Extinction-Cladogenesis (DEC) models, all estimated by the BioGeoBEARS R package (Matzke, 2016). As BioGeoBEARS requires an ultrametric tree of species/population, we used the BEAST chronogram from both mtDNA and ddRAD analyses, and all outgroups and specimens were pruned using Mesquite v.3.7 (Maddison & Maddison, 2021).

These models enable the exploration of different possibilities of vicariance, extinction, and dispersal. Furthermore, by incorporating a founder-event parameter (+J) into the analysis, we attained clastogenic dispersal outside of the parental areas (referred to as jump speciation). The *Ligula* species complex was divided into five biogeographic realms: (1) Palearctic, (2) Afrotropical, (3) Indomalayan, (4) Australasian and (5) Nearctic. Inferred ancestral ranges were allowed to occupy up to five areas, thereby facilitating clear estimations. Likelihood-based measures of three biogeographic models along with their modified J types were ranked according to the AIC method to evaluate model quality.

We applied phylogenetic trees from both mtDNA and ddRAD data to reconstruct ancestral host fish affiliations using a Bayesian framework implemented in sMap v.1.0.7 (Bianchini and Sánchez-Baracaldo, 2021). The software uses stochastic mapping analysis to reconstruct the evolutionary history of discrete characters at the branch and node of the tree. In the present study, *Ligula* plerocercoids were collected from 41 fish species representing 18 genera in four orders (Cypriniformes, Osmeriformes, Gobiiformes and Galaxiiformes). Character states were defined based on the host taxonomy at the order level. Therefore, we defined four different states as follows: 1) lineage China infects Osmeriformes, 2) *L. pavlovskii* was associated with Gobiiformes, 3) lineages of Australia and New Zealand infect Gobiiformes and Galaxiiformes, and 4) lineages of Ethiopia, EAST African Rift (EAR) and lineages A and B were associated with Cypriniformes (Supplementary Material, section 8).

### Genome-wide analysis of hybridization and introgression

Treemix v.1.12 (Pickrell & Pritchard, 2012) was applied to examine gene flow among populations. Utilizing allelic frequency data, the analysis generated an ML tree, followed by inferring the historical migratory events between populations. We assumed n migration events and the calculation of each migration event was done separately. We used the third ddRAD data set and grouped it into 11 populations based on phylogenetic analysis. The R package ‘optM’ (Fitak, 2021) was then used to optimize the number of migration events according to Evanno et al. (2005) (Supplementary Material, section 9). Visualization of the graph results was accomplished by the plot function (plotting_funcs.R) in R. We also employed the ABBA-BABA (D-statistics) and F statistics tests to further investigate patterns of gene flow among populations (Patterson et al., 2012). This analysis provides a basic yet comprehensive framework for deciphering deviation from a strictly bifurcating phylogeny. ABBA-BABA statistics were calculated in DSUITE v.0.4 (Malinsky et al., 2021) using the DTRIOS command for 139 specimens from 11 lineages and two specimens from *D. latus* as outgroup (Supplementary Material, section 9).

### Demographic history

The Extended Bayesian Skyline Plot (EBSP) (Heled & Drummond, 2008) was used to separately analyse the demographic changes for each lineage based on the Cyt *b* and COI data for inference of past population dynamics over time. EBSP analyses were implemented in BEAST v.2 under the GTR + G + I (invariant sites) model and a strict molecular clock. We employed the same mutation rate as in the divergence time analysis. We used the plotEBSP R function to draw the skyline plot (Heled, 2015). The *X*-axes indicate time reported in units of thousands of years before present (BP), while the *Y*-axes display mean effective population size (*Ne*) in millions of individuals divided by generation time plotted on a log scale. Moreover, The stairway plot2 (Liu & Fu, 2020) was used for estimating contemporary effective population sizes and changes in effective population size of the *Ligula* species complex over time. SambaR R package was used to convert the second ddRAD data (unlinked SNPs) file into folded SFS files (see details in Supplementary Material, section 10).

## Results

### Summary statistics

We compared the genetic diversity of all parasite populations based on the mtDNA (162 specimens) and ddRAD (139 specimens) data. The concatenated mtDNA matrix with 162 *Ligula* individuals and 801 bp aligned positions (404 bp Cyt *b* and 397 bp COI) was characterized by 535 monomorphic sites and 246 polymorphic sites (23 singletons and 223 parsimony informative sites). We also added 62 ND1 sequences (892 bp) to the concatenated data set for all phylogenetic analyses based on ML and BI methods. Estimates of nucleotide diversity for the two mitochondrial genes ranged from 0.0025 to 0.0196 and the haplotype diversity was between 0.61 and 1. The mean observed and expected heterozygosity ranged between 0.0005–0.044 and 0.0023–0.104, respectively. Additionally, populations from China, Australia and New Zealand, and Central and South Africa (CSA) demonstrated the lowest levels of haplotype diversity and observed and expected diversities (except *L. alternans*). The *Ligula* population from Ethiopia (12.8) showed the highest number of private alleles; however, populations in Lineage B, Australia and New Zealand, *L. alternans* and CSA had the fewest private alleles (5.8) (Table 1).

**Table 1.**
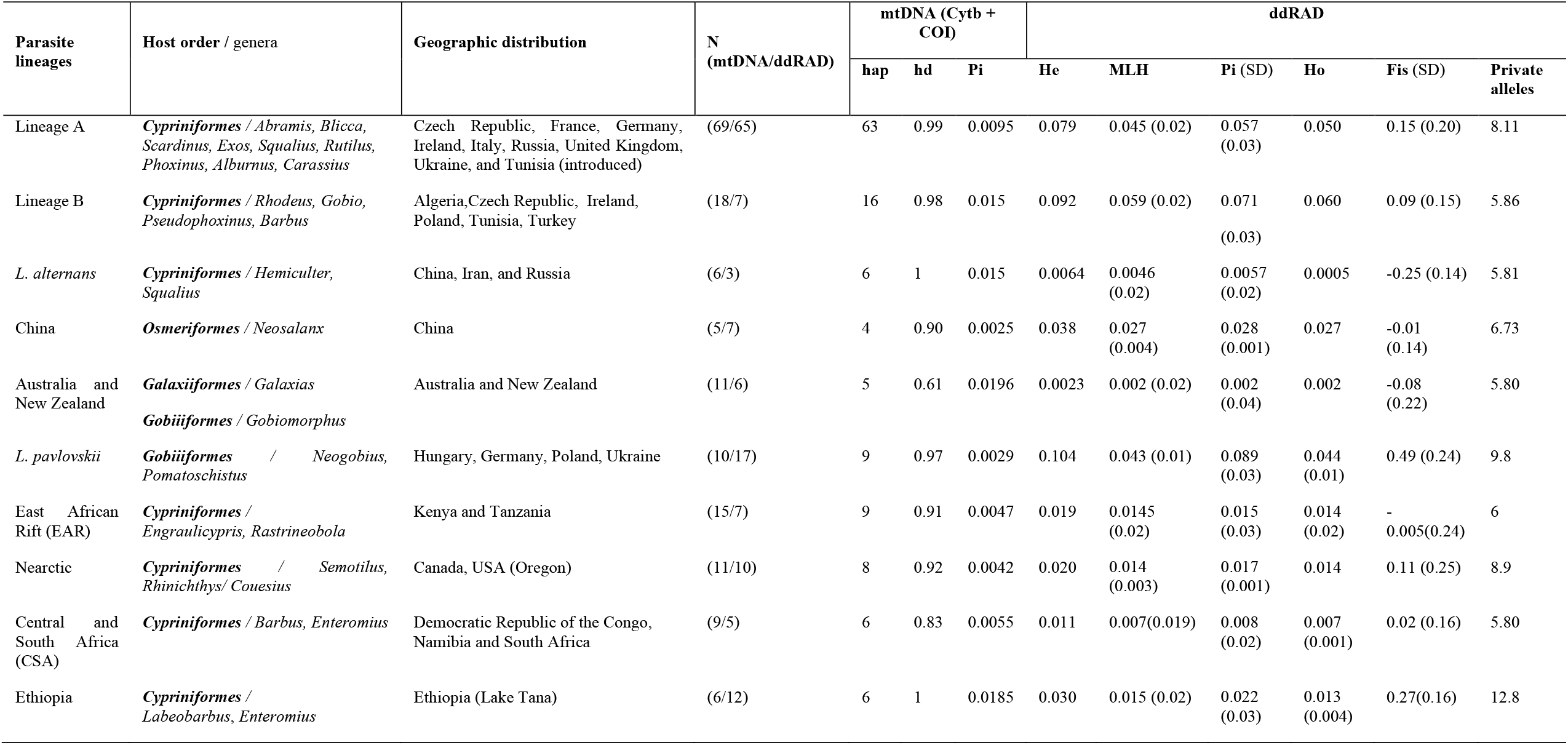
Genetic characteristics of different parasite populations based on the mtDNA and ddRAD data sets. hap: Number of mitochondrial haplotypes, hd: Mitochondrial haplotype diversity, Pi: Mitochondrial nucleotide diversity, He: Expected heterozygosity, MLH: Multi-locus heterozygosity, Ho: Observed heterozygosity, Fis: Inbreeding coefficient.

| Parasite lineages | Host order / genera | Geographic distribution | N (mtDNA/ddRAD) | mtDNA (Cytb + COI) |  |  | ddRAD |  |  |  |  |  |
| --- | --- | --- | --- | --- | --- | --- | --- | --- | --- | --- | --- | --- |
|  |  |  |  | hap | hd | Pi | He | MLH | Pi (SD) | Ho | Fis (SD) | Private alleles |
| Lineage A | *Cypriniformes* / *Abramis*, *Blicca*, *Scardinius*, *Exos*, *Squalius*, *Rutilus*, *Phoxinus*, *Alburnus*, *Carassius* | Czech Republic, France, Germany, Ireland, Italy, Russia, United Kingdom, Ukraine, and Tunisia (introduced) | (69/65) | 63 | 0.99 | 0.0095 | 0.079 | 0.045 (0.02) | 0.057 (0.03) | 0.050 | 0.15 (0.20) | 8.11 |
| Lineage B | *Cypriniformes* / *Rhodeus*, *Gobio*, *Pseudophoxinus*, *Barbus* | Algeria, Czech Republic, Ireland, Poland, Tunisia, Turkey | (18/7) | 16 | 0.98 | 0.015 | 0.092 | 0.059 (0.02) | 0.071 (0.03) | 0.060 | 0.09 (0.15) | 5.86 |
| *L. alternans* | *Cypriniformes* / *Hemiculter*, *Squalius* | China, Iran, and Russia | (6/3) | 6 | 1 | 0.015 | 0.0064 | 0.0046 (0.02) | 0.0057 (0.02) | 0.0005 | -0.25 (0.14) | 5.81 |
| China | *Osmeriformes* / *Neosalanx* | China | (5/7) | 4 | 0.90 | 0.0025 | 0.038 | 0.027 (0.004) | 0.028 (0.001) | 0.027 | -0.01 (0.14) | 6.73 |
| Australia and New Zealand | *Galaxiiformes* / *Galaxias*<br>*Gobiiformes* / *Gobiomorphus* | Australia and New Zealand | (11/6) | 5 | 0.61 | 0.0196 | 0.0023 | 0.002 (0.02) | 0.002 (0.04) | 0.002 | -0.08 (0.22) | 5.80 |
| *L. pavlovskii* | *Gobiiformes* / *Neogobius*, *Pomatoschistus* | Hungary, Germany, Poland, Ukraine | (10/17) | 9 | 0.97 | 0.0029 | 0.104 | 0.043 (0.01) | 0.089 (0.03) | 0.044 (0.01) | 0.49 (0.24) | 9.8 |
| East African Rift (EAR) | *Cypriniformes* / *Engraulicypris*, *Rastrineobola* | Kenya and Tanzania | (15/7) | 9 | 0.91 | 0.0047 | 0.019 | 0.0145 (0.02) | 0.015 (0.03) | 0.014 (0.02) | - 0.005(0.24) | 6 |
| Nearctic | *Cypriniformes* / *Semotilus*, *Rhinichthys*/ *Couesius* | Canada, USA (Oregon) | (11/10) | 8 | 0.92 | 0.0042 | 0.020 | 0.014 (0.003) | 0.017 (0.001) | 0.014 | 0.11 (0.25) | 8.9 |
| Central and South Africa (CSA) | *Cypriniformes* / *Barbus*, *Enteromius* | Democratic Republic of the Congo, Namibia and South Africa | (9/5) | 6 | 0.83 | 0.0055 | 0.011 | 0.007(0.019) | 0.008 (0.02) | 0.007 (0.001) | 0.02 (0.16) | 5.80 |
| Ethiopia | *Cypriniformes* / *Labeobarbus*, *Enteromius* | Ethiopia (Lake Tana) | (6/12) | 6 | 1 | 0.0185 | 0.030 | 0.015 (0.02) | 0.022 (0.03) | 0.013 (0.004) | 0.27(0.16) | 12.8 |

### Phylogenetic relationships and species trees of *Ligula* spp

BI and ML analyses of the three mitochondrial genes (1693 bp) generated consistent trees with a considerably well-supported phylogenetic structure (Fig. 2A). Similarly, the reconstructed phylogenies based on ddRAD data (4471 SNP) showed identical topologies in the BI and ML methods (Fig. 2B). The phylogenetic structure revealed maximum support for all lineages in both mtDNA and ddRAD data sets. In all reconstructions, the *Ligula* population from Ethiopia was strongly supported by high posterior probability (pp=1) and bootstrap support (BS=100) as the basal lineage to all the tapeworms. Parasite populations from the Nearctic separated from other lineages with maximal statistical robustness. All phylogenetic trees from mtDNA and ddRAD data disclosed a sister group relationship between populations from EAR (Tanzania and Kenya) and CSA populations. The phylogenetic relationships of the lineages derived from mtDNA data were in agreement with the phylogeny of the ddRAD data (except China and *L. pavlovskii* lineages). Within mtDNA, *L. pavlovskii* diverged from Lineage A and *L. alternans* with high statistical support. However, *L. pavlovskii* was distinctly separated from Lineage A, China, *L. alternans*, Lineage B, Australia and New Zealand. To sum up, our results added five new distinct lineages (*L. pavlovskii*, Australia, New Zealand, EAR and CSA) to the six previously defined *Ligula* lineages (Bouzid et al., 2008; Štefka et al., 2009). The SVDquartet species trees recovered the same topology as the ML and BI analyses but with variable posterior probability and bootstrap values (Fig. 2).

**Fig. 2.**
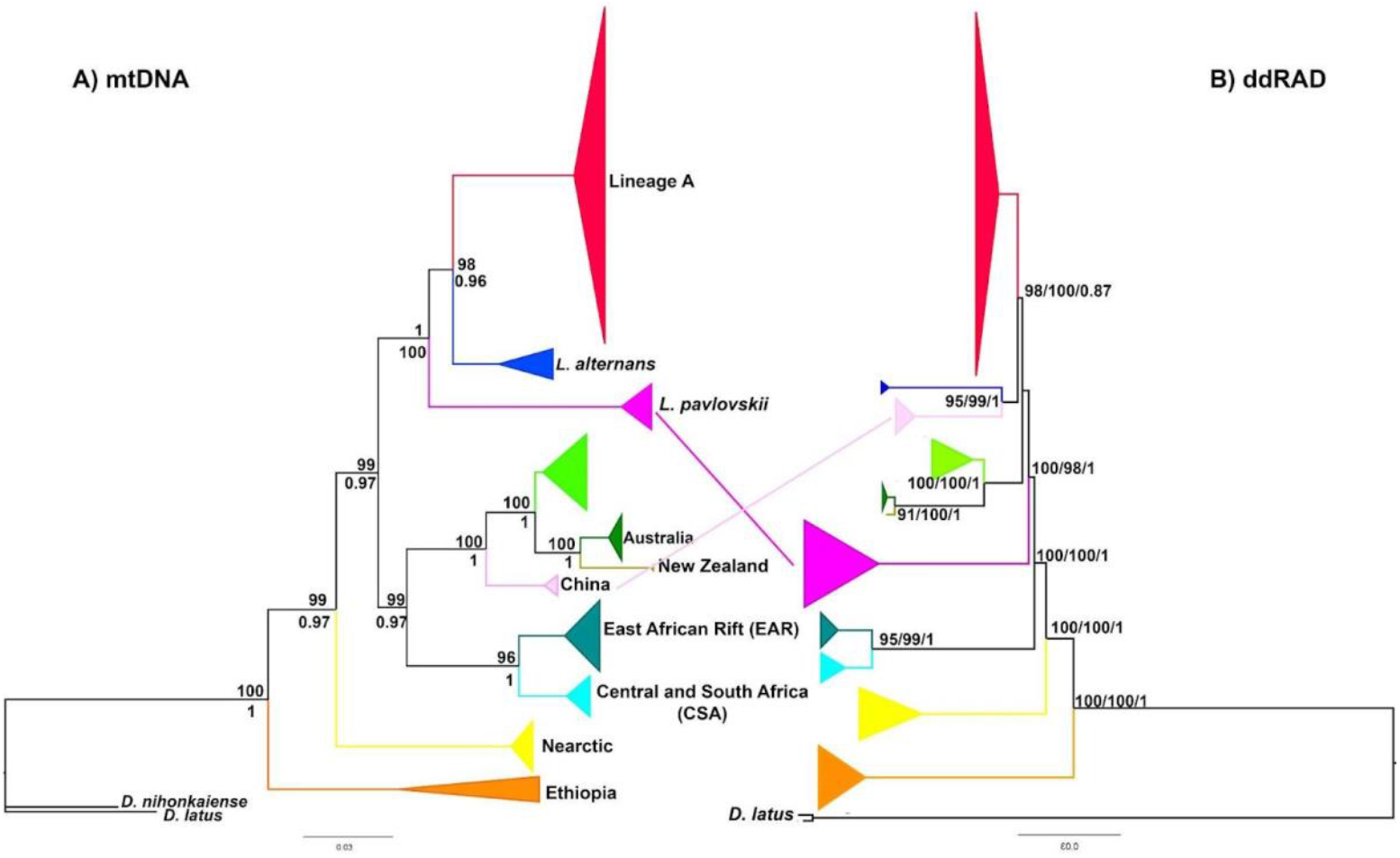
Phylogenetic relationships of the *Ligula* species complex. A) Bayesian tree reconstructed using the concatenated mitochondrial genes (Cyt *b*, COI and ND1) using two outgroups (*D. latus* and *D. nihonkaiensis*). Lineage positions and branching patterns are concordant with the ML tree. Nodal supports at each node represent support values of BI (above branches) and ML (below branches). B) Reconstructed phylogenetic tree based on ddRAD. Bootstrap support values and Bayesian posterior probabilities are given at the nodes (RAxML/svdquartets/ExaBayes). Diagonal lines represent incongruences between the mtDNA and ddRAD topologies. Topology congruence tests demonstrated significant differences between the mtDNA and ddRAD trees (P < 0.05).

### Population genetic structure and host usage

A total of 133 haplotypes were found based on the Cyt *b* and COI sequences in 162 individuals. The mitochondrial haplotype network based on TCS analysis with a 95% parsimony connection limit revealed the same 11 distinct haplogroups as in the phylogenetic analysis (Fig. 3A, connection limit =12 mutations). Individual haplogroups showed a high level of host specificity. Although many lineages were retrieved from Cypriniformes, the genera (and families) of the fish rarely overlapped between lineages even in areas of sympatry (such as Europe). Haplogroup A is widely distributed in Europe, Iran and Russia and included *Ligula* populations from Cypriniformes (*A. brama, R. rutilus, B. bjoerkna, S. erythrophthalmus, S. cephalus, A. alburnus, P. phoxinus* and *C. carassius*). The *L. alternans* haplogroup was connected to haplogroup A with 39 mutational steps and comprised parasite populations from Iran, China, and far east Russia, which were associated with *Hemiculter* (Cypriniformes: Xenocyprididae) and *Neosalanx* (Osmeriformes: Salangidae). Haplogroup B was linked to Australia and China with 30 and 36 mutational steps, respectively. Furthermore, haplotypes from Australia and New Zealand diverged from each other with 41 mutational steps and included parasite populations from Galaxiiformes (*Galaxias*) and Gobiiformes (*Gobiomorphus*). Lineage B and the haplogroup in China were found in Cypriniformes (*Rhodeus, Gobio* and *Pseudophoxinus*) and Osmeriformes (*Neosalanx*), respectively. The haplogroup in EAR comprised parasite populations of *E. sardella* that diverged from the haplogroup in CSA (*E. anoplus* and *E. paludinosus*) by 22 mutational steps. The *Ligula* populations from Nearctic Cypriniformes (*S. atromaculatus* from Canada and *R. osculus* from Oregon) split from the other lineages by 55 and 120 mutational steps. Furthermore, haplogroups of *L. pavlovskii* from Gobiiformes (*N. fluviatilis, N. melanostomus, A. fluviatilis, P. minus* and *P. microps*) and Ethiopia from Cypriniformes (*L. intermedius, L. tsanenis, L. brevicephalus* and *E. humilis*) separated from each other by 110 mutational steps (Fig. 3A). Contrary to the mitochondrial haplotype network, the phylogenetic network based on ddRAD data clustered Australia and New Zealand together, revealing 10 well-supported clusters for the *Ligula* species complex (Fig. 3B). Ethiopia was the first lineage branching at the base. EAR and CSA split from each other with 98% bootstrap support value, diverging from *L. pavlovskii* and Ethiopia with 100% bootstrap support value. Similarly, Australia and New Zealand differed from Lineage B. Also, Lineage A demonstrated a divergence from all parasite populations with strong bootstrap value (100%).

**Fig. 3.**
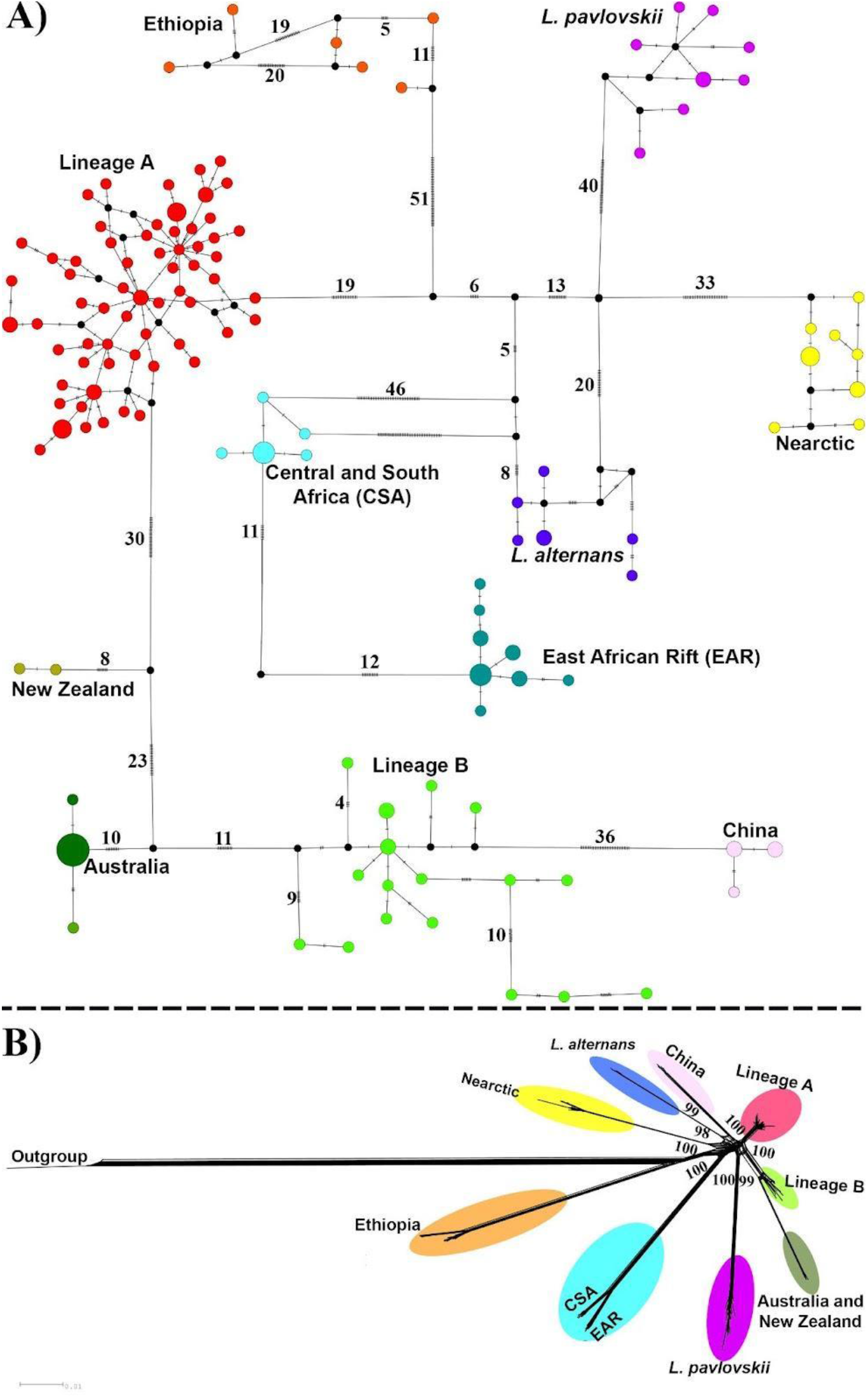
Population relationships. A) Median-joining network showing relationships among the *Ligula* species complex, reconstructed based on the concatenated Cyt *b* and COI sequences (801 bp). Circle sizes are proportional to haplotype frequencies. Black circles indicate unsampled or extinct haplotypes. Mutational steps are displayed by numbers and dashed symbols along each line. B) Phylogenetic network with bootstrap support values for the ddRAD data showing 10 distinct clusters in the *Ligula* species complex. *D. latus* was used as an outgroup.

Population structure analyses showed the three new clusters (*L. pavlovskii*, EAR, CSA) were clearly separated from the five previously identified lineages (Lineage A, *L. alternans*, Australia, Lineage B and China) (Fig. 4). Cross-validation of the admixture analysis revealed an optimal number of K=10 clusters (Fig S2). All lineages were separated from each other, although some admixture was observed between EAR and CSA (Fig. 4A). PCA patterns were mainly analogous to the clustering recognised by admixture. The first two PCs represented 22.6% and 18.1% of the total genetic variation, separated into 10 groups. EAR was close to CSA and had an overlap on both PC1 and PC2 axes. The third PC (16%) clearly discriminated Lineage A, Lineage B and Ethiopia from the remaining groups (Fig. 4B). Similarly, the cross-entropy curve of the spatial population genetic structure recovered a likely number of K=10 genetic clusters (Fig. S3). We spatially interpolated the ancestry coefficients projected onto a map to describe the global genetic structure in our data (Fig. 4C). The resulting spatial pattern revealed 10 components of ancestry in the sampling space.

**Fig. 4.**
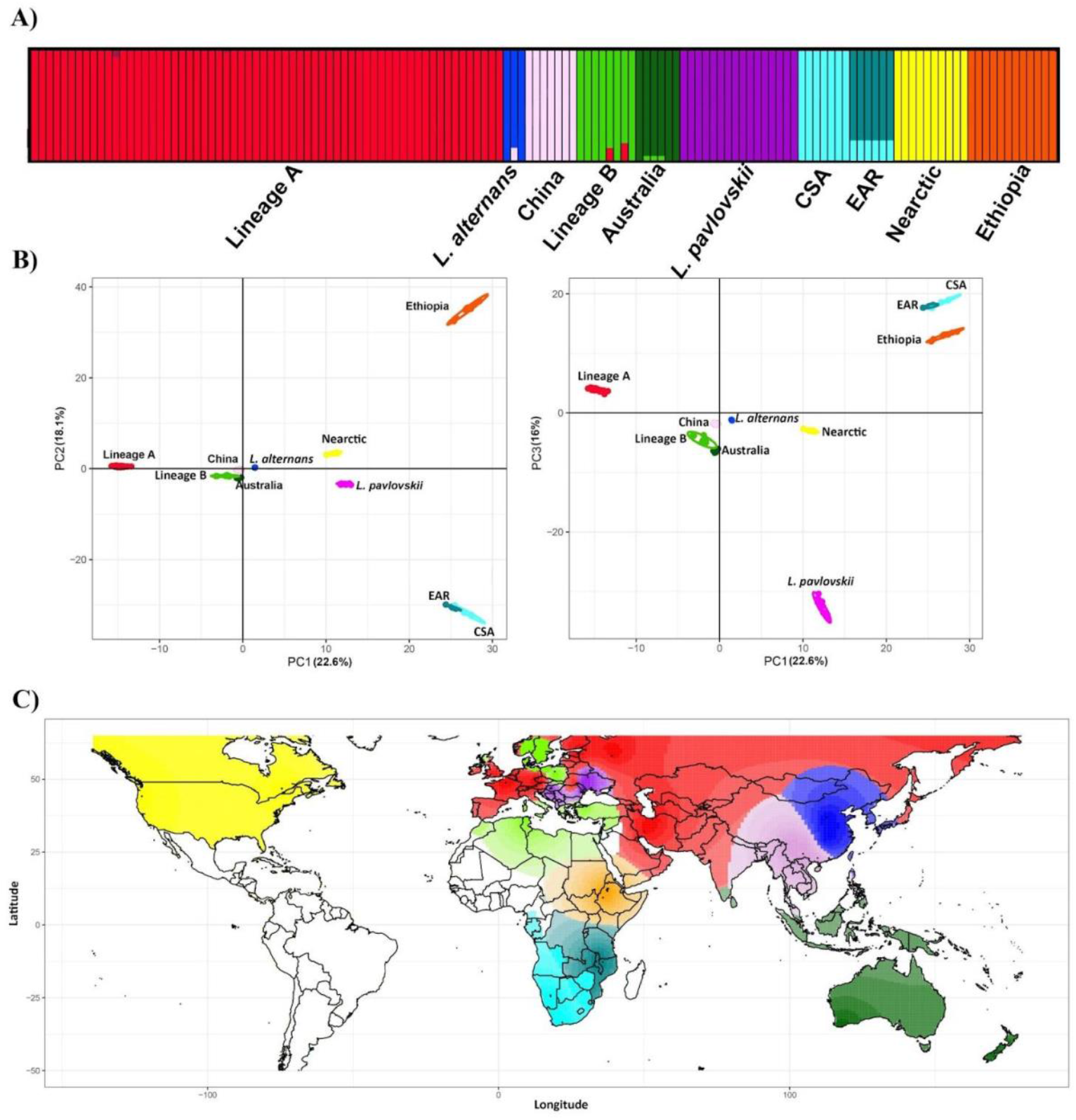
Genetic structure of the populations. A) Population structure based on the ADMIXTURE analysis. Results are shown at K = 10. Each vertical (100%) stacked column indicates an individual representing the proportions of ancestry in K-constructed ancestral populations. B) PCA of *Ligula* sp. samples with 10937 SNPs, where the first three components explain 22.6%, 18.6% and 16% of the variance, respectively. C) Interpolated spatial genetic structure according to the individual ancestry coefficients of TESS3R. The colour gradient shows the degree of difference among individuals.

The analysis of shared ancestry in the fineRADstructure confirmed the results of the cluster analysis with K=10, with significant support in the dendrogram. Although the recent co-ancestry was greatest between EAR and CSA, the two lineages formed distinct groups in the analysis. Moreover, a single sample from New Zealand showed a higher level of shared genetic background with the cluster from Australia. These two clusters had relatively higher co-ancestry with Lineage B and China. Moreover, lineages A and B exhibited a moderate common ancestry with each other, whereas the lowest level of co-ancestry sharing was discovered among Ethiopia, *L. pavlovskii*, Nearctic, EAR and CSA. This is consistent with the admixture results where a level of admixed population was observed between EAR and CSA as well as Australia and Lineage B (Fig. 5).

**Fig. 5.**
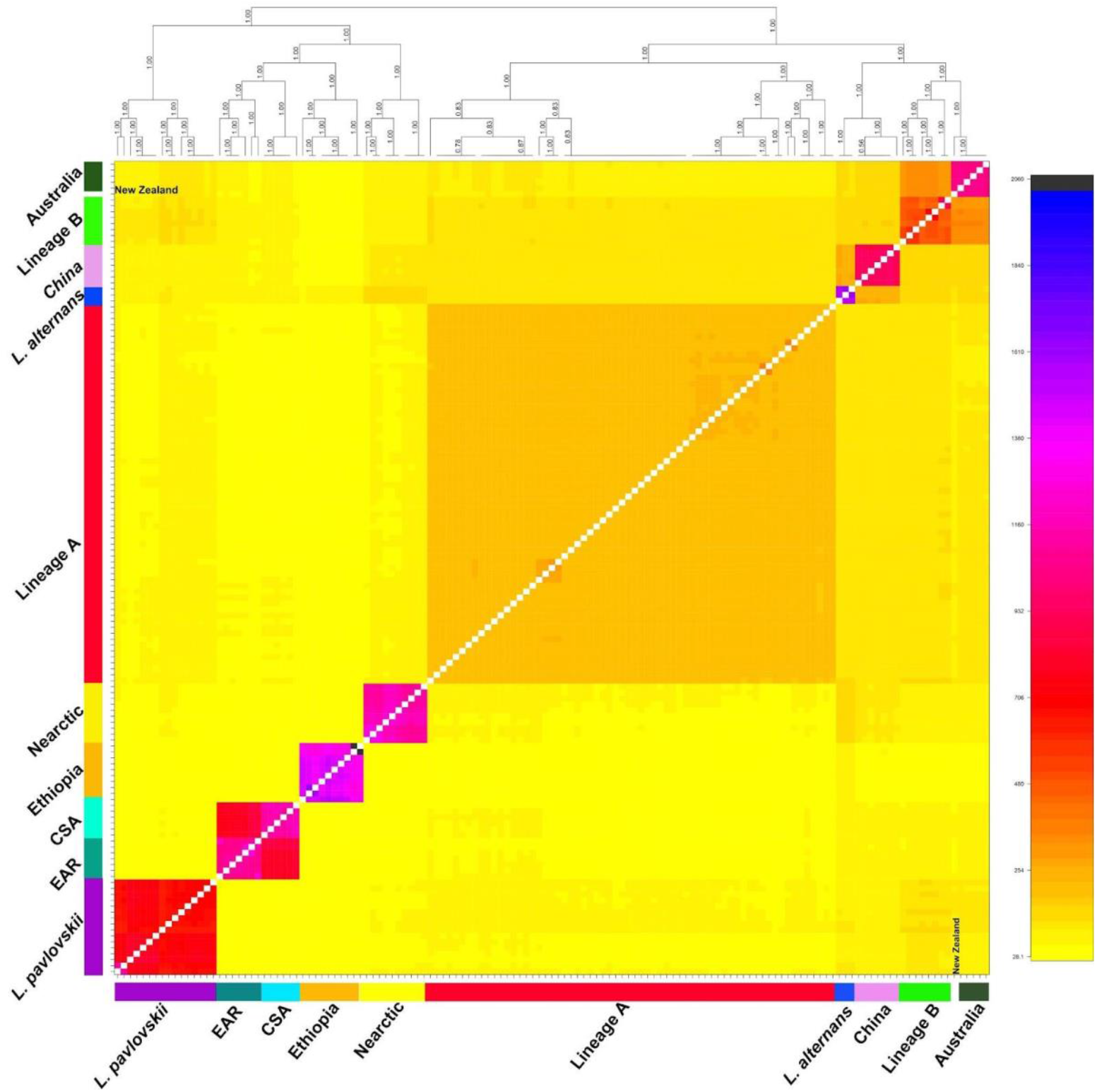
Pairwise genetic similarity among individuals indicated by FineRADStructure co-ancestry matrix. Inferred populations are clustered together in their accompanying dendrogram.

### Species delimitation analyses

The species delimitation methods (ASAP, GMYC, bPTP and BFD) produced estimates that ranged from 10 to 15 species (GMYC and bPTP) (Fig. 6). ASAP analysis with the best score delimited 10 and 12 species based on Cyt *b* (lowest score = 2.5), COI (lowest score = 2.5) and ND1 (lowest score = 4). In the ND1 fragment, out of the 10 top-scoring models, three showed distance thresholds of 5.7%, 1.4% and 2.7% with seven, 13 and 12 species. The third was selected as the most likely initial estimate of species diversity. Tree-based analyses identified the same composition and number of species. The results of the BFD hypothesis testing are shown in Fig. 4, Fig. S6 and Table S4. The ddRAD data set favoured 10 candidate species based on marginal likelihood (3820.21, Fig. S4) and BF (32.5). Overall, all delimitation analyses determined Lineage A, *L. alternans, L. pavlovskii*, China and Nearctic as different species. However, GMYC and bPTP over-split Lineage B and Ethiopia into three candidate species. In addition, all the methods (except ASAP based on Cyt *b* and BFD) distinguished a species boundary between Australia and New Zealand.

**Fig. 6.**
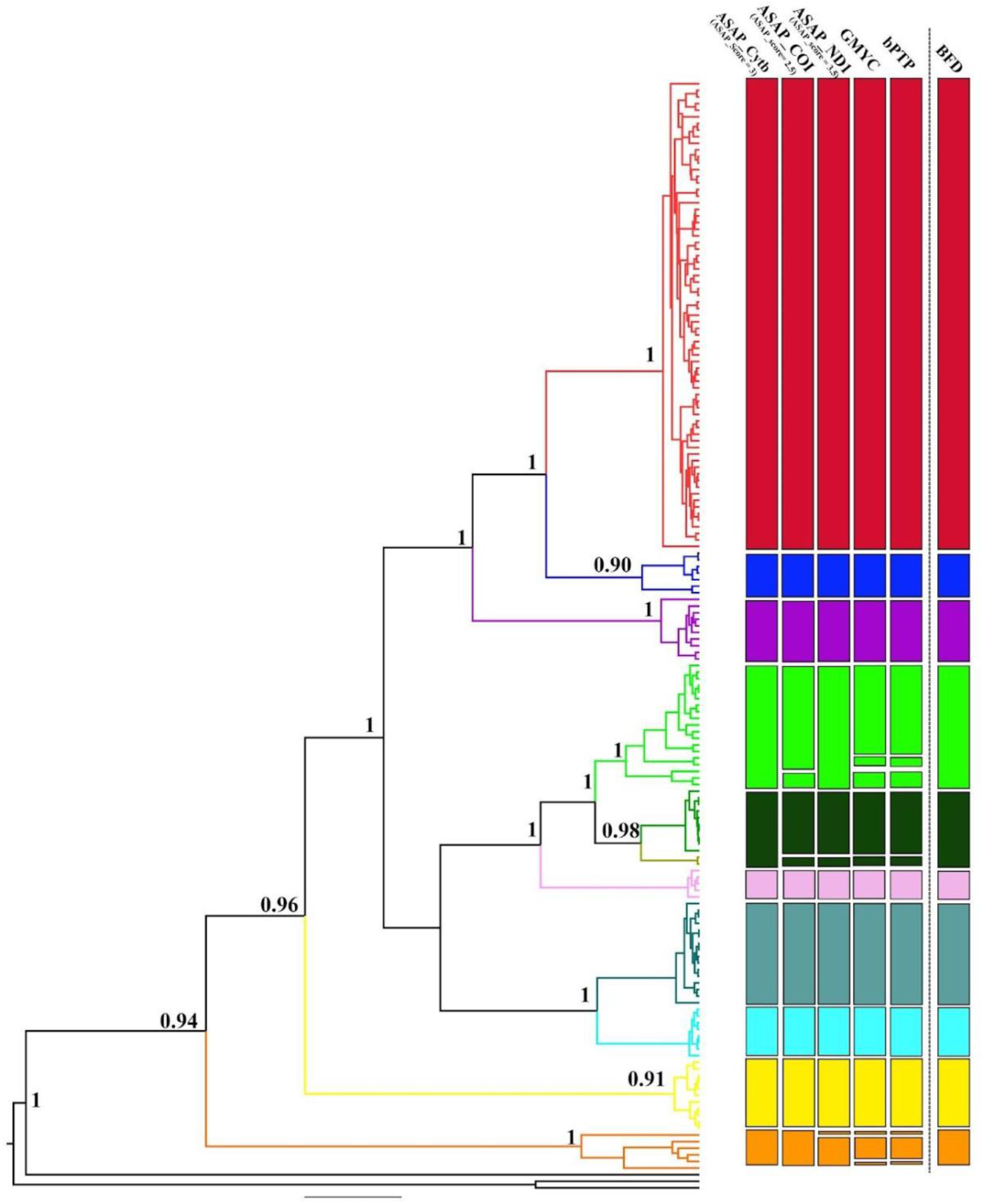
Multi-approach species delimitation of *Ligula* species complex. Results of the mtDNA method (ASAP, GMYC, bPTP) are compared to the results of genome-wide SNP methods (BFD). Coloured bars depict results from different methods, indicating congruence among them.

### Divergence time and phylogeographic structure

Estimate of the divergence times using mitochondrial genes was consistent with the time-calibrated tree of ddRAD, strongly supporting the basal cladogenesis between lineages of the *Ligula* species complex in late Miocene and early Pliocene (mtDNA= 4.99 Mya, 95% highest posterior density intervals (HPD): 4.76–5.27 Mya and ddRAD =5.05 Mya, HPD: 4.56–5.49), with the Ethiopian population emerging as the basal lineage to all *Ligula* populations (Fig. 7). Subsequent cladogenetic events dating to mid-Pliocene at approximately 3.35–4.01 Mya (mtDNA=4.01 Mya, HPD= 4.87–3.10 Mya and ddRAD=3.35 Mya, HPD= 3.04–3.78 Mya) isolated the ancestor of Nearctic lineage from other populations. Both mtDNA and ddRAD results revealed a sister relationship between EAR and CSA which diverged from each other in early Pleistocene at 1.01–1.14 Mya (mtDNA= 1.14 Mya, HPD: 0.71–1.41 Mya and ddRAD =1.01 Mya, HPD: 0.66–1.29 Mya). Furthermore, both dated trees strongly supported a sister relationship between Australia and New Zealand which diverged from Lineage B at ∼1.12– 1.23 Mya (mtDNA= 1.12 Mya, 95% HPD: 0.78–1.81 Mya and ddRAD =1.23 Mya, HPD: 1.11– 1.49 Mya). In mtDNA, *L. pavlovskii* diverged from *L. alternans* and Lineage A at ∼2.41 Mya (Fig. 7A, HPD: 0.78–2.96 Mya); however, ddRAD revealed that *L. pavlovskii* split from the other six lineages at ∼2.48 Mya (Fig. 7B, HPD: 2.29–2.88 Mya). Furthermore, in mtDNA, China diverged from Australia, New Zealand and Lineage B at ∼1.71 Mya (Fig. 7A, HPD: 1.11–2.17 Mya), while in ddRAD, China formed a sister relationship to *L. alternans*, diverging at ∼1.52 Mya (Fig. 7B, HPD: 1.09–1.85 Mya).

**Fig. 7.**
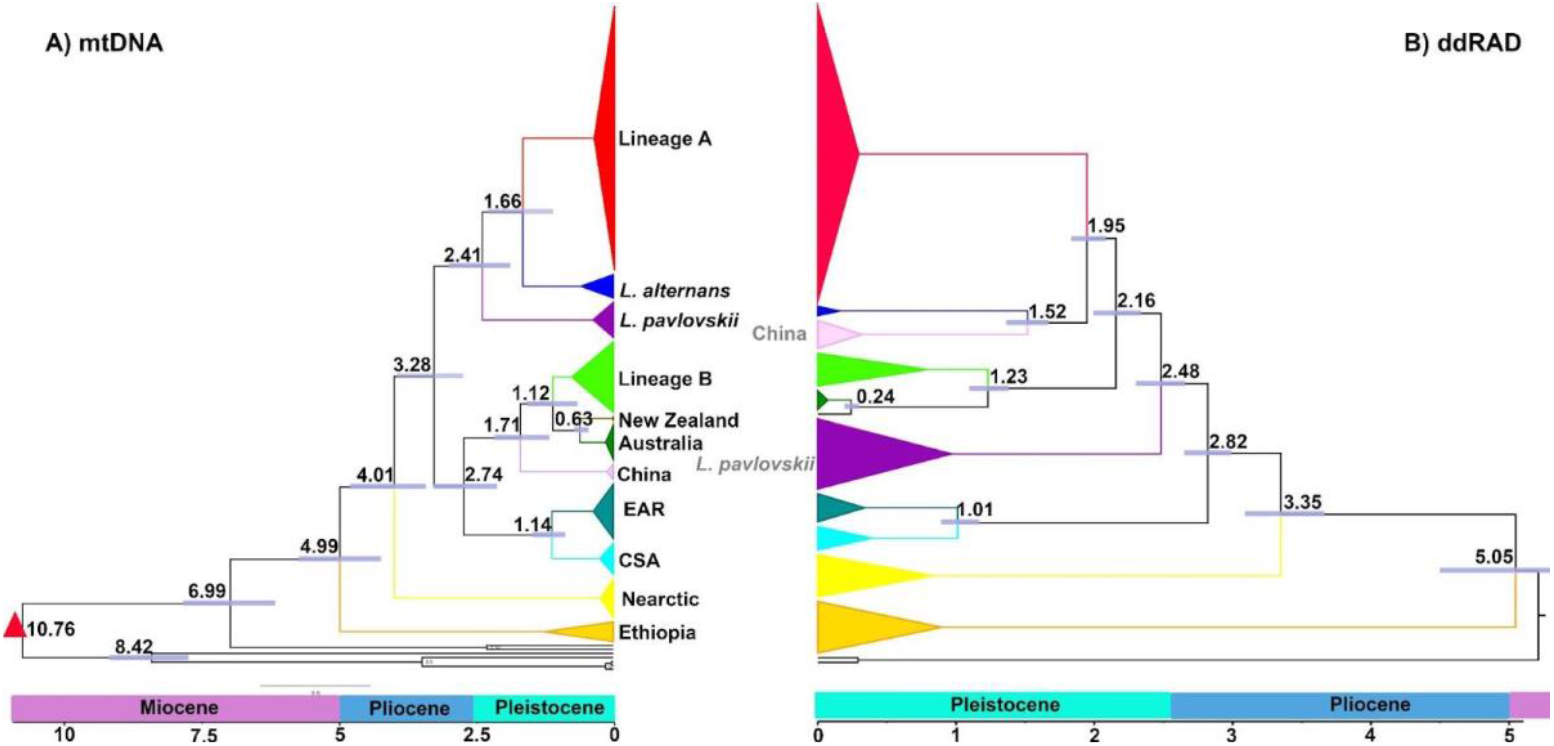
Chronograms obtained from the dating analyses. A) Concatenated data set of three mitochondrial genes (Cyt *b*, COI, and ND1) containing 165 *Ligula* sequences. Numbers on each branch indicate divergence times. Calibration point is displayed by a red triangle. B) Divergence time estimated from the ddRAD data set (4471 unlinked SNPs). The bars on the nodes represent 95% HPD for divergence times.

### Historical biogeography and gene flow analyses

Ancestral range estimation analyses were applied to chronograms resulting from both the mtDNA and ddRAD data sets (Fig. 8A). A comparison of the six biogeographic models with the likelihood ratio test and AIC indicated that DIVALIKE+J was the top fitting model for both the mtDNA and ddRAD data sets (Tables S4). The preferred hypothesis would be that the common ancestor was widespread throughout the Palearctic, Afrotropical and Nearctic biogeographic realms between the late Miocene and early Pliocene. Moreover, vicariance took place between Nearctic and Palearctic in mid-Pliocene. With respect to anagenetic or gradual speciation processes, our parameters indicate that range contraction made an equal contribution as did range expansion in both data sets (Table S4). The ancestor of EAR and CSA emerged through a rare jump dispersal event from the Palearctic to the Afrotropical in late Pliocene. In addition, the China lineage was formed as a result of founder event speciation between Palearctic and Indomalaya in mid-Pleistocene. Finally, Australia and New Zealand were genetically isolated from their ancestor populations in the Palearctic (Fig 8A).

**Fig. 8.**
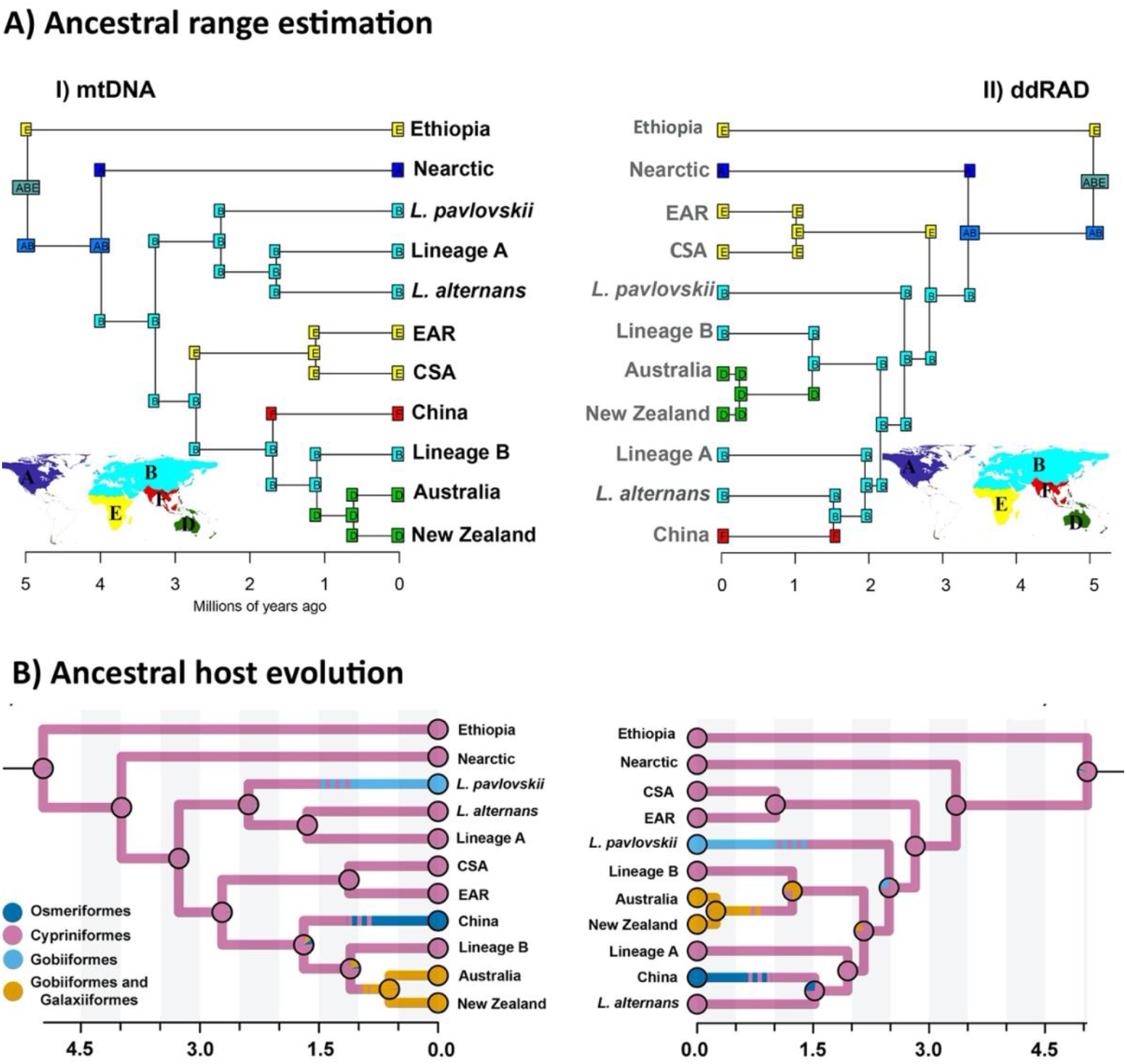
Ancestral range and host estimations. A) Ancestral range estimation for the tapeworm chronograms in BioGeoBEARS under the DIVALIKE+J model. Squares at nodes indicate the highest ML probability in the area prior to the sudden speciation event, while squares on branches refer to the states of descendant lineages immediately after the speciation event. Squares showing more than one letter represent ancestral areas with more than one biogeographical area. B) Bayesian stochastic character map displaying the ancestral host evolution based on mtDNA (I) and ddRAD (II). Posterior probabilities for each node in different states are depicted by pie charts.

Of the three-character models evaluated using sMap, the ER model was the best supported (BIC, Table S5) for both mtDNA and ddRAD data sets. The Bayesian stochastic character mapping of ancestral host fish associations demonstrated an ancient affiliation with cypriniforms at the root of all *Ligula* lineages (Fig 8B). In both data sets, mean marginal posterior estimates showed that cypriniforms contributed as the most recent common ancestral host for the parasite with 81.4% and 82.3% in mtDNA and ddRAD, respectively. Also, the most recent common ancestor (MRCA) of *Ligula* had the highest host switching from cypriniforms to the other fish orders. Furthermore, *Ligula*’s MRCA had the second highest average time spent (mtDNA =9.1%, ddRAD= 12.1%) in Gobiiformes and Galaxiiformes, and the lowest average time spent in Osmeriformes (Table S4, Supporting Material, mtDNA=5.5% and ddRAD=6.1%). A host shift from cypriniforms towards Gobiiformes and Galaxiiformes was discovered in both data sets for the MRCA of Lineage B, Australia and New Zealand. In addition, we identified the occurrence of several host shifts for the MRCA of China and *L. alternans*. On the other hand, the mtDNA chronogram revealed the MRCA of *L. pavlovskii* experienced three host shifts from Cypriniformes to Gobiiformes and Osmeriformes, while ddRAD stochastic map detected a host shift between Cypriniformes and Gobiiformes (Fig. 8B).

### Genome-wide hybridization and introgression

The estimated species tree from Treemix recovered similar topology (Fig. 9A) and relative branch lengths to the phylogenetic trees. The simplest model that described the data was the model that assumed three migrations. No significant migration events were retrieved by more complex models (Fig S4). The results revealed gene flow from Nearctic into *L. alternans* and China populations. Moreover, two gene flow events were detected from Lineage B into China and *L. pavlovskii*. This finding shows how introgression affects phylogenetic relationships in closely related species. To determine possible introgression events, we performed ABBA-BABA tests for 10 trios of closely related phylogroups. D statistics for 47 of the trios were significantly different from zero (Z > 3, p-adjusted <.000919) and the values ranged between 0.244 and 0.492 (Table S6). Based on the sliding window analysis, SNPs demonstrating the ABBA-BABA patterns for each trio were distributed across 25–29 different contigs (Table S3). Nevertheless, it is possible that a single gene flow event leads to multiple elevated f4-ratio and D results. When we calculated a branch-specific statistic fb(C) that partially disentangled the correlated f4-ratio results, we found that only two out of the eight branches exhibited significant excess sharing of derived alleles at least with one other lineage (Fig. 10B). An internal branch that represented Lineage A and China exhibited an excess sharing of derived alleles with Australia, New Zealand, *L. pavlovskii* and Lineage B. Another fb(C) signal involves an internal branch as the common ancestor of Lineage B with *L. pavlovskii*, EAR and CSA (Fig. 9B). These results could also be attributed to incomplete lineage sorting. It must be noted that even a single introgression event may result in significant fb(C) values in multiple related phylogroups (Malinsky et al., 2018).

**Fig. 9.**
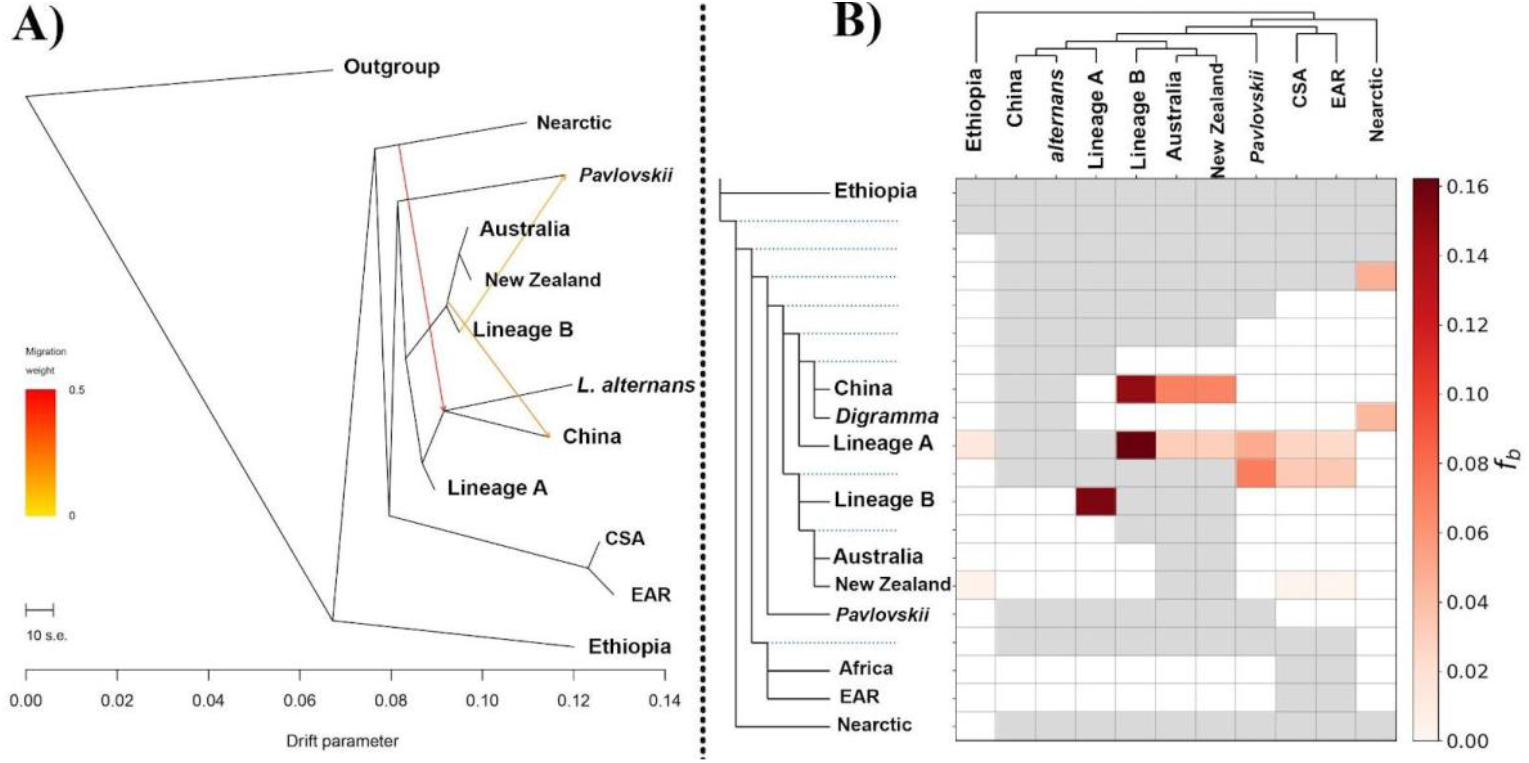
Patterns of historical gene flow A) ML tree from Treemix, representing the relationships between the 11 *Ligula* lineages. The scale bar displays 10 times the average standard error for entries estimated in the covariance matrix. Migration arrows (events) are coloured based on their weight. The fraction of ancestry from the migration edge is referred to as migration weight. B) Results of genome-wide *f*-branch among the 11 *Ligula* lineages. Columns represent tips in the tree topology and rows indicate tree nodes. Cells display fb statistics between nodes (rows) and tips (columns). When comparisons cannot be made, the cells are left empty and grey. F-branch (fb(C)) statistics in our data suggest excess allele sharing among tips of the phylogenetic tree (representing lineages which are arranged horizontally at the top) and between each tip and node in the tree (arranged vertically on the left).

**Fig. 10.**
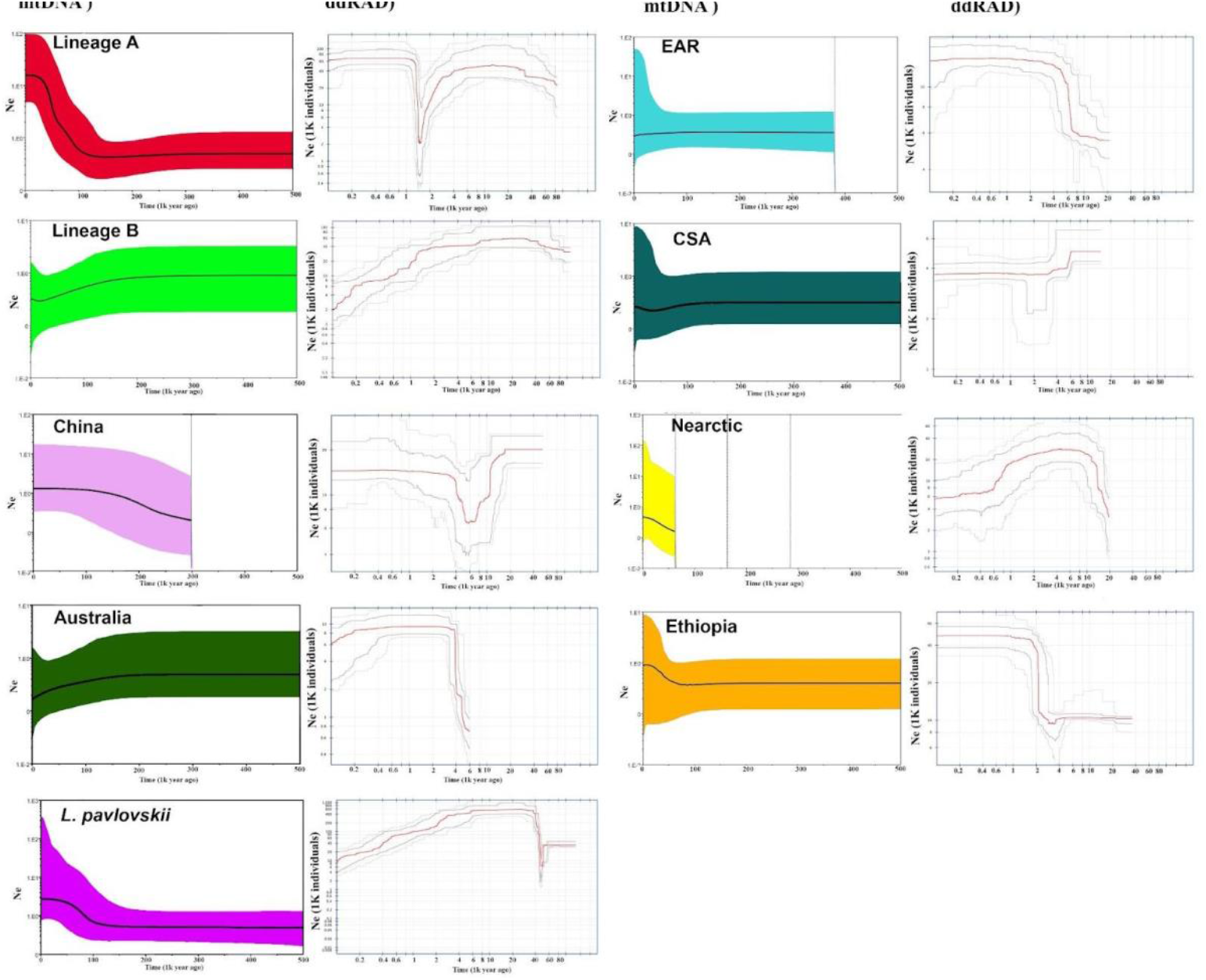
Demographic history analyses. The plot on the left is EBSP based on two mtDNA genes (Cyt *b* and COI) for each main lineage. Bold line represents the median and the coloured area indicates 95% confidence interval. On the right, the stairway plot shows calculated changes in effective population size (Ne) for each lineage over time estimated from site frequency spectrum data. The grey lines above and below the red line (mean estimation) indicate 75% (dark) and 95% (light) confidence intervals.

### Historical and contemporary demographics

The demographic history based on two mtDNA genes (Cyt *b* and COI) and ddRAD data sets is presented for nine parasite lineages in Fig. 10. Two lineages (*L. alternans* and New Zealand) were not plotted due to their low effective sample sizes (ESS<200) and low sample sizes. EBSP based on the mtDNA suggested that demographic expansion took place at approximately 100 kya (thousand years ago) for lineages A and B (Fig. 10 A and B). Moreover, the stairway plot based on the ddRAD data showed a clear sign of ancestral bottleneck between 1.5 kya and 2 kya followed by a range expansion for these lineages. China showed a population expansion after experiencing a bottleneck between 4 kya and 8 kya. Australia demonstrated a trend indicative of recent population shrinkage. Our mtDNA results uncovered a rapid population expansion of approximately 100 kya for *L. pavlovskii*; however, ddRAD revealed a bottleneck at ~50 kya with a decreased pattern from 6 kya onwards. A stable population size is indicated for EAR and CSA during the last 3 kya. In addition, a recent population reduction and a strong signal of demographic expansion were detected for the Nearctic and Ethiopian lineages.

## Discussion

Host specificity, an inherent feature of parasitic and symbiotic organisms, can be a product of co-diversification (Engelstädter & Fortuna, 2019), but it is often the result of a more convoluted evolutionary history including host switches and dispersal events (Hoberg & Brooks, 2008; Matthews et al., 2022). The latter should be peculiar to parasites where the host lineages are much older than the parasite lineages, such as the *L. intestinalis* species complex. Using our example of a speciating parasite lineage with a global distribution, we demonstrated how host specificity of individual *Ligula* lineages was formed by different historical events involving long-distance dispersals, host switches and occasional introgression. Our analyses based on a dense SNP matrix and mtDNA data provided a clear picture of the cryptic diversity within the *L. intestinalis* species complex and a robust time-calibrated phylogeography. By analysing biogeographic and demographic patterns of individual lineages, we revealed patterns of the parasite’s speciation and dispersal throughout its distribution range. Below, we discuss the results focusing on individual factors contributing to the parasite’s diversification.

### Phylogenetic relationships and occasional mito-nuclear discordance

*Ligula* phylogenies based on ddRAD and mtDNA data revealed 11 distinct lineages and showed a generally well-supported phylogenetic structure for the *Ligula* species complex. We characterized several new lineages from previously unexplored areas (CSA, EAR) or lineages that were not analysed in the global context before (*L. pavlovskii*, New Zealand; see Lagrue et al., 2018; Vitál et al., 2021). In addition, several differences in the phylogeny of the group were recovered compared to previous studies based on fewer loci and less material (Bouzid et al., 2008; Štefka et al., 2009). Ethiopia formed a basal lineage to other populations in our analysis, contrary to the findings of the previous study (Bouzid et al., 2008) where Nearctic samples (from Canada and Mexico) were at the basal position. Our dataset also allowed for a more fine scale distinction of the lineages previously clustered with Lineage B (Bouzid et al., 2008).

Only two cases of discordance between mitochondrial and nuclear-ddRAD phylogenies were found, with *L. pavlovskii* and China showing different topologies in each data set. To exclude a possibility that the incongruence is only due to the lack of signal in the smaller mtDNA dataset, we performed a congruence test between mtDNA and ddRAD topologies and found the topology differences were significant (AU and SH tests). Similar mito-nuclear phylogenetic discordances have also been reported in several other metazoan taxa (e.g. Bouzid et al., 2008; Shaw, 2002; Weigand et al., 2017), often credited to the molecular signature of selection (Morales et al., 2015), incomplete lineage sorting (Degnan & Rosenberg, 2009) or introgressive hybridization (Bisconti et al., 2018). Our analyses of historical gene flow indicated past introgression events between multiple lineages (see the discussion below). Thus, we suggest introgressions as the mechanism responsible for the emergence of mito-nuclear discordances for China and *L. pavlovskii* lineages.

### Species delimitation reveals extensive cryptic diversity in *L. intestinalis* s.l. populations

The taxonomy of *Ligula* tapeworms has always been complicated due to the general uniformity of their plerocercoids and adults (Dubinina, 1980), resulting in a lack of suitable morphological features to determine species boundaries using traditional methods, except for an already known trait, the duplicated reproductive complexes in *L. alternans* (Luo et al., 2003). Indeed, numerous species of *Ligula* have been proposed to accommodate plerocercoids from different fish hosts (see Dubinina, 1980 for their extensive list), but almost all these taxa have been synonymised with *L. intestinalis*. This species was then considered as a euryxenous parasite of cyprinoid fishes, maturing in several fish-eating birds (Kuchta & Scholz, 2017; Li et al., 2018; Petkevičiūtė, 1992).

Here, we used a range of species delimitation analyses to identify the putative species in the *Ligula* complex. The results based on ddRAD data revealed a slightly lower number of candidate species than the non-recombining mtDNA. Our findings also indicated that although different methods of species delimitation produced highly consistent species boundaries, they were not entirely similar. Carstens et al. (2013) argued that such incongruence either signifies a difference in the power of one or more of the methods to detect cryptic lineages or that assumptions of the species delimitation approach have been violated. They suggested that species delimitation inferences should be drawn cautiously in order to avoid delimiting entities that fail to accurately represent evolutionary lineages. Therefore, taking the conservative approach, we identified at least 10 putative species for the *Ligula* species complex as follows: (1) *L. intestinalis* Lineage A; (2) *L. alternans* (syn. *Digramma alternans*); (3) *L. intestinalis* Lineage B; (4) *Ligula* sp. 1 from Australia and New Zealand (possibly two species lineages); (5) *L. pavlovskii*; (6) *Ligula* sp. 2 from China; (7) *Ligula* sp. 3 from EAR; (8) *Ligula* sp. 4 from CSA (9); *Ligula* sp. 5 from the Nearctic; and (10) *Ligula* sp. 6 from Ethiopia. In subsequent analyses, we contrasted these evolutionarily significant lineages with their host and geographic spectra and demographic histories.

### Dated phylogeography reveals historical dispersal and vicariance due to climatic oscillations and host switching

Climatic fluctuations have shaped the patterns of divergence in a multitude of species (Hewitt, 2001). In particular, quaternary glaciation cycles and climatic oscillations had a major impact on both free-living and parasitic species, leading to the extinction of several parasites (Hoberg et al., 2012; Li et al., 2018) and influencing their evolution and distribution (e.g. Galbreath et al., 2020; Nazarizadeh et al., 2022b). Based on our mtDNA and ddRAD molecular dating, the ancestor of *Ligula* is estimated to have appeared between the late Miocene and early Pliocene (Fig. 7), and was found to parasitise one of the early diverging cyprinid species (*Barbus*, of the subfamily Barbinae). The divergence time suggests that the parasite complex did not follow the evolution of its intermediate host as the divergence of Cypriniformes had already occurred during the late Jurassic (154 Mya) (Tao et al., 2019). Moreover, the historical biogeography model revealed that the divergence of parasite populations from Afrotropical, Nearctic and Palearctic realms was recovered as two vicariant events (Fig. 8). This allopatric speciation could be associated with the isolation of migratory waterbirds due to the biogeographical realms becoming isolated by major barriers. According to our results, the first vicariance occurred between Afrotropical, Palearctic and Nearctic regions at approximately 5 Mya (Fig. 8) when desertification increased and the decline of global temperatures accelerated, resulting in the development of the Northern Hemisphere ice sheets and the onset of the ice ages. Moreover, the considerable expansion of the Sahara Desert had by then impeded trans-Saharan migration and long-distance dispersal for several migratory waterbirds (Bruderer & Salewski, 2008).

*Ligula* plerocercoids mature very quickly into adults in bird guts following ingestion of an infected fish (Loot et al., 2001). The adult phase lasts up to five days (Dubinina, 1980), during which the eggs can be dispersed. Consequently, a lack of suitable freshwater lake habitats within a few days flight of the final host represents a strong barrier against gene flow of the parasite, as documented by the lack of gene flow in Lineage A between Europe and a recently introduced population in North Africa (Bouzid et al., 2013; Štefka et al., 2009). Thus, aridification of the environment accompanied by reduced or ceased migration of the definitive host likely caused the major vicariance mechanisms in the evolution of *Ligula*.

The second vicariant event took place between the Nearctic and Palearctic during the late-Pliocene when the Nearctic lineage diverged from the other populations (Fig. 8). This lineage was recovered from the Nearctic leuciscids of two subfamilies (*Semotilus* - Plagopterinae, and *Rhinichthys* - Pogonichthyinae) which are thought to have originated in ancient Europe and dispersed to North America during the Late Cretaceous to Paleocene (see Imoto et al., 2013). In contrast to the intermediate hosts which had evolved a long time before *Ligula*’s speciation, changes in the behaviour of the definitive hosts could have accelerated parasite’s diversification from Pliocene to Pleistocene (Hirase et al., 2016). It has been proposed that glaciations possibly shifted the migratory routes and migratory behaviour of several North American bird species (Zink & Gardner, 2017). In addition, ecological competition and physical isolating barriers between the Palearctic and Nearctic during the late Pliocene or early Pleistocene were determining factors (Naughton, 2003). On the other hand, the results of TreeMix showed historical gene flow between the population from the Nearctic and the MRCA of lineages of China and *L. alternans* (Fig. 9A). This introgression likely occurred after the second vicariant event when both the Nearctic and Palearctic regions were connected through the Bering Land Bridge, which allowed for a passage of many species between the two realms (Beaudoin & Reintjes, 1950).

Founder event speciation is a form of allopatric speciation in which a small number of individuals establishes a new population beyond the existing range of the main population (Matzke, 2014). The results of the historical biogeographic model revealed that a jump dispersal event took place from the Palearctic to the Afrotropical in the early Pleistocene, forming the MRCA of EAR and CSA (Fig. 8). In line with the present study, several fossil records of waterbirds have been discovered from the early Pleistocene in EAR, the Upper Pliocene in Ethiopia as well as Miocene deposits in Kenya (Brodkorb & Mourer-Chauviré, 1982; Dyke & Walker, 2008; Prassack et al., 2018). Our results do not support the theory of *Ligula* colonization of Africa in recent decades (Gutiérrez & Hoole, 2021; Kihedu et al., 2001). Instead, we suggest that the parasite had already colonized sub-Saharan Africa in the early Pleistocene. Moreover, the dated tree based on both mtDNA and ddRAD data indicated that parasite populations from EAR and CSA diverged from each other in the late Pleistocene (Fig. 7). EAR lineage showed host specificity to *Engraulicypris sardella* and *Rastrineobola argentea*, whereas CSA lineage to barbels (*Enteromius anoplus, E. paludinosus, E. trimaculates* and *Barbus neefi*). Whilst barbels are a common host for *Ligula* in many regions (Barson & Marshall, 2003; Emaminew et al., 2014) and represent one of its ancestral hosts, a switch to the hosts endemic to the EAR region could have played a role in the differentiation of the EAR lineage.

Our biogeographic analyses also suggested the occurrence of sympatric speciation in *Ligula*. The historical biogeographic model revealed that three lineages (Lineage A, Lineage B and *L. pavlovskii*) were formed as a result of two sympatric speciation events in the western Palearctic. It also revealed that the MRCA of these lineages were historically distributed in the Palearctic (Fig. 8A and B). Moreover, some samples from these lineages were found in the same waterbodies. Lineages A and B were found in two different groups of cyprinid fish and *L. pavlovskii* was discovered in gobies, i.e., members of the distantly related fish order Gobiiformes. Similarly, Bouzid et al. (2008) revealed that the Euro-Mediterranean lineages (lineages A and B) live in sympatry and infect the same definitive host. They also suggested that reproductive isolation formed a significant genetic barrier between the two lineages. Moreover, a local genetic population study on *Ligula* provides preliminary evidence of a host-related ecological differentiation in Lineage A at an intra-lineage level (Nazarizadeh et al., 2022a). Reproductive isolation in sympatry is mainly a consequence of the interaction between various pre- and post-zygotic barriers (Nosil, 2012). The proximate mechanisms of differentiation in sympatric lineages of *Ligula* are yet to be explored; however, for instance, Henrich & Kalbe (2016) indicated that postzygotic ecological selection is more likely the cause of restricted gene flow between *Schistocephalus solidus* and *S. pungitii*, sister species of tapeworms related to *Ligula*.

The phenomena of geographical dispersal and host-switching are common in parasites at different spatial and temporal scales (Hoberg & Brooks, 2008). Cyclical and episodic climatic changes are predicted to influence ecological release and the response of populations to the relaxation of isolation mechanisms that increase host-switching potential (e.g. Brooks et al., 2006; Hoberg et al., 2002). Through host-switching, a small proportion of a species moves into a new geographical area, which may be followed by speciation through peripheral isolation and addition of a new species to the parasite’s host range (Huyse et al., 2005). The results of divergence times and historical biogeography confirmed that the latest “jumping speciation” in *Ligula* occurred from the Palearctic to Indomalayan and Australian realms in the mid-Pleistocene (approximately 2.0 to 1.2 mya). The host switch from Cypriniformes to native freshwater fish (Osmeriformes in China, Galaxiiformes and Gobiiformes in Australia and New Zealand) led to “peripheral isolate speciation” (Frey, 1993), a pattern previously reported in several cestode species (Brooks, 1989; Zarlenga et al., 2006).

### Genome-wide hybridization and introgression

Results of the TreeMix introgression and the f-branch tests were highly consistent, with the f-branch statistics revealing a higher number of gene flow events. Introgression results suggested the presence of several ancestral ghost lineages between Ethiopia and other lineages (Fig. 10B). Ghost lineages, indicative of the existence of unsampled or extinct lineages in a species, significantly influence gene flow detection (Tricou et al., 2022). TreeMix detects introgression events based on the known lineages and branches in a tree and does not consider ghost lineages (Tricou et al., 2022).

Historical and contemporary patterns of hybridization have been documented for many tapeworms (Bello et al., 2021; Easton et al., 2020; Landeryou et al., 2022). Henrich and Kalbe (2016) confirmed natural hybridization between *S. solidus* and *S. pungitii* from both allopatric and sympatric populations. Our results detected historical hybridization in both sympatric (Lineage A, Lineage B and *L. pavlovskii*) and allopatric lineages (Nearctic, China, Lineage A, Australia and New Zealand). In addition, we discovered significant positive D-statistic values, indicating an excess of derived allele sharing between China and Lineage B, in line with the TreeMix migration edge (Fig. 10B). We suggest that historical introgressive hybridization might have enabled *Ligula* to infect a broader range of host species (Osmeriformes, Gobiiformes and Galaxiiformes), similar to adaptation of hybrids of nonparasitic taxa to novel ecological conditions where their parents could not persist (Thaenkham et al., 2022). However, future testing of specific questions related to adaptive introgression will require extended sampling and a well-resolved and annotated genome assembly.

### Demographic history

Demographic analyses based on mtDNA and ddRAD data showed mutually consistent results for most lineages, with SNP based results providing a more recent picture (Fig. 11). EBSP showed Lineage A and Lineage B experienced rapid range expansions (Fig. 11A) during the Last Interglacial Period (LIG; Otto-Bliesner et al., 2006). Moreover, during the Medieval Warm Period (MWP; Crowley & Lowery, 2000), these lineages exhibited rapid expansions following a bottleneck event (Fig. 11B). Although these very recent events are not straightforward to explain without more comprehensive data and analyses, they fit into the general picture of a volatile population history expected for parasitic taxa, involving bottlenecks and expansions following host-switches as well as expansions due to post-glacial host range expansions. Periods of geographical and host range expansions allow parasites to explore their capacity for generalism, while periods of isolation force parasites to specialise on specific hosts. This specialisation will modify the “sloppy fitness space” (Agosta & Klemens, 2008; Araujo et al., 2015) of the parasite due to the parasite adapting to local environments.

Bottlenecks found in *Ligula* lineages, particularly those associated with lineage formation, could be initiated by host switches followed by population expansion in case of a successful specialisation. During a host switch, it is likely that the host immune system can decrease overall parasite load, while at the same time it selects for more immune-resistant parasite’s genotypes (Levin et al., 1999). *Ligula* interacts with the immune and metabolic systems of the fish host intensively, leading to the castration of the host and redirecting energy resources towards the parasite’s growth (Yoneva et al., 2015). Future analysis of coding genetic differences and dating their origin, ideally using whole genome data, could allow testing the hypotheses of host-switch associated bottlenecks.

## Acknowledgements

we are grateful to Anna Mácová and Roman Hrdlička for their help with the fieldwork. The authors also wish to thank Eva Čisovská, Košice, Slovakia for providing *D. latus* samples to generate outgroup sequences. The research was supported by a grant from the Czech Science Agency (no. GA19-04676S). Computational resources were supplied by the project “e-Infrastruktura CZ” (e-INFRA CZ LM2018140) supported by the Ministry of Education, Youth and Sports of the Czech Republic.

## Data Accessibility Statement*

Raw sequence reads are deposited in the SRA (BioProject XXX). Individual genotype data are available on DataDryad (XXXX). MtDNA data are deposited to NCBI Nucleotide Database (XXXX). *Data will be made available prior to publication of the manuscript in a peer-reviewed journal.

## Benefit-Sharing Statement

All authors took part in field-collecting samples for the research via a collaborative effort.

## Author Contributions

MNaz. and JŠ defined the research objective and drafted the manuscript. MNov., PK and MNaz. performed laboratory analyses. MNaz. analysed the data under the supervision of JŠ and ET. All authors took part in sample collecting and read and approved the final version of the manuscript.

