## Supplemental Information for "Historical dispersal and host-switching formed the evolutionary history of a globally distributed multi-host parasite - the *Ligula intestinalis* species complex"

### Table of Contents:


| | |
|---|---|
| Table S1a and S1b | Supplementary Tables.xlsx |
| Supplementary material, section1 | Page 2 |
| Supplementary material, section2 | Page 2 |
| Supplementary material, section3 | Page 3 |
| Supplementary material, section4 | Page 3 |
| Supplementary material, section5 | Page 4 |
| Supplementary material, section6 | Page 5 |
| Supplementary material, section7 | Page 6 |
| Supplementary material, section8 | Page 6 |
| Supplementary material, section9 | Page 6 |
| Supplementary material, section10 | Page 7 |
| Table S2 | Page 8 |
| Table S3 | Page 8 |
| Table S4 | Page 9 |
| Table S5 | Page 9 |
| Table S6 | Page 10 |
| Figure S1 | Page 11 |
| Figure S2 | Page 12 |
| Figure S3 | Page 12 |
| Figure S4 | Page 13 |
| Figure S5 | Page 14 |

### Section 1

#### Polymerase chain reaction (PCR) and mitochondrial sequencing

Each gene fragment was amplified in a 12 ul PCR containing 6.25ul 2x concentrated PPP Master Mix (Top-Bio, CZ), 1 ul of extracted DNA and 10 pM of forward and reverse primers. The amplification and sequencing primers used in this study are listed in Table S3. The protocol for amplification of Cyt *b* (405-900 bp) and COI genes (396 bp) consisted of an initial denaturation step for 15 min at 94 °C, followed by 30 cycles of 30 s denaturation at 94 °C, annealing for 45 s at 50 °C, and extension for 45 s at 72 °C. The last elongation step was carried out for 10 min at 72 °C. The same procedure was applied for the ND1 gene (892 bp), except for the annealing temperature which was set at 55 °C. VWR ExoCleanUp FAST PCR reagent (VWR, USA) was used to enzymatically clean up PCR products following the manufacturer's instructions. Purified PCRs were Sanger sequenced in a commercial laboratory (Seqme, CZ) using PCR primers.

### Section 2

#### ddRAD library preparation

ddRAD libraries were prepared following the Peterson et al. (2012) protocol with modifications specified below. Briefly, genomic DNA was digested at 37 °C for 2 h using the restriction enzymes NspI and MluCI (NEB). Complete digestion was verified on 1,5% agarose gel. The Ligation step for 30ul reactions contained 0.2uM P1-flex adapter, 1uM P2-flex adapter, 1X T4 ligase buffer, 400U T4 DNA ligase, ddH2O and standardizing amount of digested DNA per sample in a range of 76-100ng for each individual library. Reaction conditions were: 22°C for 60 minutes following by enzyme heat inactivation at 65°C for 10 minutes at ramp rate -1°C/min. A total of 48 uniquely barcoded samples in each library were pooled into one tube, vortexed and distributed equally into 18 strip tubes. Two steps of magnetic beads purification were carried out. First, all samples were diluted by 10 ul ddH2O and combined into two tubes before magnetic plate incubation. Then, the resulting DNA was second cleaned and diluted by 16 ul ddH2O. The Contents of both tubes were combined together before final magnetic plate incubation. Finally, pooled samples were verified with Qubit 2.0 Fluorometer and size selected in a range of 275-325 bp (Pippin Prep, Sage Science). Finally, PCR reactions were performed combining 6.25ng of size-selected sample, PCR primers 1 and 2 at final concentration 0.2uM each, the recommended amount of 5X-HF buffer, dNTPs, ddH2O and Phusion polymerase, and run on 12 cycles. Quality and concentrations of final libraries were assessed using agarose gel electrophoresis and a Qubit 2.0 fluorimeter, respectively. Libraries were sequenced using Illumina Novaseq 150 PE reads in a commercial laboratory (Novogene, UK).

#### Section 3

### SNP filtering of assembled ddRAD data

ddRAD assembly was aided by mapping the reads against a draft genome of *L. intestinalis* obtained in the frame of a concurrent study. After assembly in the Stacks (Rochette et al., 2019), we applied several different filters to the population component of stacks to keep loci that were present across 90% of all populations (p), in a minimum of 80% of individuals from each population (r=0.80) where the minor allele frequency (MAF) was 0.05 and demonstrated a maximum observed heterozygosity of 80% (max_obs_het= 0.80). As regards PCA, phylogeny analyses and FST-based analyses, we discarded SNPs in linkage disequilibrium with the squared coefficient of correlation 0.5 in 50-SNP sliding windows. In addition to the post-processing step, further filtering of variants was carried out in vcftools v0.1.16 (Danecek et al., 2011), removing variants with low (<5x) and high depth of coverage (800x). Moreover, all loci with higher than 10% missingness were discarded.

#### Section 4

### Phylogenetic analyses

The best-fitting substitution models were estimated using the Bayesian Information Criterion (BIC, Table S3). Bayesian and maximum likelihood phylogenetic trees were reconstructed using the selected models in Mrbayes and IQtree, respectively. The Bayesian analysis was carried out with one cold and three heated chains (MC3) for 60 million generations and sampled every 2000th generation with a burn-in of 25%. We examined convergence in the Tracer v1.5 package (Andrew Rambaut et al., 2018), which was then visualised with the convergence diagnostic parameters from MrBayes.

Raxml-ng (Kozlov et al., 2019) was run using the GTR+G substitution model to perform 50 ML tree searches with 25 random and 25 parsimony-based starting trees. 1000 nonparametric bootstrap replicates were generated based on the best tree search. Convergence was checked post-hoc using the --bsconverge command in raxml-ng after changing the cutoff value to 0.05; Branch supports were computed and mapped onto the best-scoring ML tree by means of Felsenstein bootstraps (FBP) and Transfer Bootstrap Expectation (TBE) (Lemoine et al., 2018). We used four coupled chains in ExaBayes (Aberer et al., 2014, all with three heated chains) to run two Metropolis-coupling replicates for 1000000 MCMC generations, sampling every 500th generation. The postProcParam tool was used to assess the estimated sample size (ESS) of all parameters and branch lengths, which was confirmed to be adequately sampled using Tracer v.1.6 (Aberer et al. 2014). Eventually, a majority-rule consensus tree was computed using the 'consense' tool from the Exabayes package (burn-in: 25%). Species trees were constructed using SVDquartets (Chifman and Kubatko, 2014) in PAUP v.4.0a147 (Swofford & Sullivan, 2003). Subsampling was performed at 10,000 quartets. The sample quartets were assembled in a species tree with a variant of the FM Quartet as recommended by the developers of SVDquartets (Chifman & Kubatko, 2014). To inspect the incongruence between the topologies obtained from mtDNA and ddRAD data sets, we tested the fit of a constrained SNP topology on the mtDNA dataset using Shimodaira-Hasegawa (SH), Approximately unbiased (AU), and Kishino and Hasegawa (KH) implemented in PAUP.

### Section 5

### Population genetic structure analyses

the AdmixPiPe (Mussmann et al., 2020) was used with a default thin parameter (t=100) and ran 20 ADMIXTURE replicates for each value of K from 1 – 20 using the second VCF file described in material and methods. The optimal values of K were calculated in ADMIXTURE based on the lowest of cross-validation error (CV) across replicate runs (Alexander & Lange, 2011, Fig. S2). CLUMPK server (http://clumpak.tau.ac.il/) was used to plot clustering individuals into populations.

The python script Stacks2fneRAD.py in the fneRADstructure package was used to calculate the distribution of alleles and SNPs per locus and of the missing data per individual. A maximum number of 10 SNP per locus (13291 loci) and a maximum 25% individual missingness (139 individual) were considered to convert the haplotype file into RADpianter format. Considering that ddRAD is susceptible to batch effect as a result of small deviations among libraries at the size selection step, we assessed whether missing data were driving any library-based structure. The fineRADstructure pipeline was run using default settings, but we raised the number of burn-in iterations to 200000, along with 1000000 iterations, and sampling every 1000. Subsequently, we calculated convergence by 1) assigning individuals to populations for multiple independent runs, 2) tracing plots for the MCMC output of the parameter values to guarantee convergence on the same Bayesian posterior distributions, and 3) generating effective parameter sample sizes (> 100) by running each chain long enough. "FinestructureLibrary.R" function in fineRADstructure was used to plot the co-ancestry heatmap (Malinsky et al., 2018).

**Section 6**

**Species delimitation analyses**

To generate an ultrametric gene tree for the analysis, we reconstructed a phylogenetic tree based on the concatenated alignment (Cyt b, COI, and ND1) using BEAST v.1.8.2 (Drummond et al., 2012), under the assumption of uncorrelated lognormal relaxed molecular clock. The analyses were performed under the constant population size coalescent as the tree prior and Ucld mean prior was set to exponential distribution with mean 10 and initial value 1. We executed analyses in two independent runs with 100 million generations, sampling every 5000. We used Tracer v.1.6 for assessing convergence and adequacy of posterior samples (ESS > 200). The first 20-25 % of trees were discarded, followed by combining the rest of trees using LogCombiner v 1.8.2. We subsequently used mean heights for annotation to compute a maximal clade credibility (MCC) tree by TreeAnnotator utility v.1.8.2 (Bouckaert et al., 2014). The GMYC approach (Pons et al., 2006) is based on discerning stochastic birth-death processes (essentially a pure-birth Yule process) between species from neutral coalescent intraspecific processes by analyzing the timing of branching events (successively summarized as combination of independent Poisson models) in single gene trees. Input prerequisites need a well-sampled, well-estimated, ultrametric single neutral locus tree which optimally characterizes the true species genealogy in absence of population structure and population size fluctuation. We used the P2C2M.GMYC r package (Fonseca et al., 2021) to test whether the data has a good fit to the GMYC model. The GMYC approach with a single threshold (GMYC) was carried out in RStudio v. 4.2 (RStudio Team, 2020.) using the Splits R package. The ultrametric tree from the Beast analyses was used for both analysis. Using the function 'multi2di', we avoided having branches of length 0. Contrary to GMYC, bPTP (Zhang et al., 2013) only needs a simple phylogenetic tree rather than an ultrametric tree. Therefore, we used the ML phylogenetic tree reconstructed in the IQTREE. The analyses were run in the bPTP servers available at http://species.h-its.org/ and http://mptp.h-its.org/#/tree. For the genetic distance-based approaches (ASAP, Puillandre et al., 2021), each gene data set was treated independently as a single partition to determine the possible putative species. The ASAP analysis only employs pairwise genetic distances to sort specimens into putative species without an a priori species hypothesis. It performs automatic detection of substantial variations between and within species. We uploaded the aligned sequences of the three genes (Cyt *b*, COI, and ND1) separately to the web server https://bioinfo.mnhn.fr/abi/public/asap. The analysis was conducted based on uncorrected pairwise genetic distances and split groups below 0.01 probability.

Bayes Factor Delimitation (BFD, Leaché et al., 2014) was carried using SNAPP package (Bryant et al., 2012) in BEAST2. The unsampled mutation rates (u, v) were both fixed at 1, alpha at 1, beta at 250 and lambda at 20. A coalescence rate initially set to 10 was sampled and default values were used for all other parameters. To achieve marginal likelihood estimation for each competing model, path sampling was performed for 48 steps in total, followed by MCMC runs of 200,000 generations (sampled every 1,000 steps). We used Bayes factors to rank and compare the resulting marginal likelihood values (Kass & Raftery, 1995). To monitor the possible influence of different priors, the analyses were repeated with default options for the mutation rates (u, v), alpha, beta, lambda, and coalescence rate. Using the highest ranked model from BFD, we performed a final species tree estimation in SNAPP. MCMC was run for 1,000,000 generations, sampling every 1,000 steps. We used Tracer to confirm high ESS values and convergence, and TreeAnnotator v.1.7.5 to construct an MCC tree with a 25% burn-in.

#### Section 7

**Divergence time and phylogeographic structure**

Substitutions and clock models were unlinked with the following settings: an uncorrelated lognormal clock for Cytb and COI and a strict clock for ND1. The dynamics of speciation and extinction were modelled by a birth-death process to infer lineage-specific rate heterogeneity. We performed four independent MCMC replicates at 400 million generations, sampled every 5000 stepovers. Tracer 1.5 was applied to evaluate the appropriate amount of burn-in and effective sample sizes of all parameters (ESS>500). Additionally, LogCombiner v.1.8.2 (Andrew Rambaut & Drummond, 2015) was used to merge all four tree files after removing a range of 18-26% as burn-in. Finally, a consensus tree with divergence times was generated in TreeAnnotator v1.8.2 with the maximum clade credibility tree and posterior probabilities of 0.95. For ddRAD data set, Four replicate MCMC runs were carried out, with the tree and parameter values sampled every 10000 generations over a total length of 5 × 108 steps. We used the diagnostic tool Tracer v1.5 in assessing convergence between runs, estimating an appropriate burn-in period to be implemented, and obtaining reasonable estimates of model parameter variance by ensuring the adequacy of effective sample sizes (larger than 500). All runs were combined using LogCombiner We used FigTree v1.3.1 (A Rambaut & Drummond, 2010) to view the sampled maximum clade credibility tree.

#### Section 8

**Ancestral host reconstructions**

We used sMap (Bianchini & Sánchez-Baracaldo, 2021) to evaluate three discrete models by means of the Bayesian information criterion (BIC): (i) all rates of transition were the same (ER); (ii) all rates of transition were different (ARD); and (iii) forward and reverse rates of transitions between states were identical (i.e., symmetrical, SYM). ER was a better fit model to both mtDNA and ddRAD data than either the ARD or the SYM model (Table S4). The phylogenetic trees were normalised using the -N/--norm option. We carried out the analysis with two independent parallel MCMC runs, sampled every 10 steps over 1000 generations, a minimum number of 2000 samples were collected from four chain analyses, and 200 samples were burin-in. We checked the convergence of Bayesian analysis using the "ChainMonitor" function (ESS > 800). Stochastic mapping was visualised in TreeViewer V 2.0.1 (https://treeviewer.org/).

### Section 9

**Genome-wide analysis of hybridization and introgression**

We specified relationships between lineages by the pruned tree for BioGeoBEARS analysis. We used *D. latus* as the root to construct the ML trees. There was a variable number of tested migration events (M), ranging between 1 and 12 (number of populations+1). Following model fit quantification by all covariances for each migration event, we evaluated the robustness of the tree topology based on bootstrap replicates created from blocks of 1,000 SNPs. Patterson's D tests and f4 -ratio statistics are computed by DTRIOS, which provides information about trios of lineages with significant D values based on Z-score, calculated with a book-jackknife technique, taking linkage among sites into consideration. We left the option jknum (the number of jackknife blocks to divide the genome into) at its default value (20 blocks of 4471 SNPs), because the results were not affected by different tested values (20,30,40, 50). The resulting p-value was adjusted by applying the Benjamin-Hochberg method for multiple testing correction via false discovery rate control. To facilitate the evaluation of correlated f4- ratio statistics, the FBRANCH command implemented the calculation of f-branch statistics and its output was plotted with dtools.py. We conducted a sliding-windows analysis for the lineage trios with significantly elevated D by means of DINVESTIGATE command to determine whether the admixture signal was limited to particular contigs (Windows size =2. Step =1) (Malinsky et al., 2021).

### Section 10

#### Demographic history

The Extended Bayesian Skyline Plot was carried in BEAST 2. We ran Markov chains for 600 × $10^{7}$ generations and sampled every 10,000th generation. Log files were analysed in Tracer v.1.7 (Rambaut et al., 2018) to evaluate the convergence of the MCMC analysis with effective sample sizes (ESS larger than 200), combined with 10% burn-in removal during 'skyline' plotting.

We ran Stairway Plot2 (Liu & Fu, 2020) using the two-epoch model, with the recommended 67% of sites used for training and 200 bootstraps on the folded SFS. Singletons were excluded from the estimation to reduce genotype calling errors. We estimated the mutation rate per generation to be 2.89 × $10^{-9}$ according to our evaluation of the divergence time. The analysis is a flexible multi-epoch model derived from the composite likelihood of an observed site frequency spectrum (SFS) and is not based on parameters or predefined models (such as constant population size followed by a bottleneck) to infer demographic histories. Therefore, a more detailed demographic history can be reflected in the model.

Table S2 List of primers used for amplification and sequencing.

| Primer | Sequences | size (bp) | Source |
|---|---|---|---|
| **Cyt *b*** | | | |
| F2Dnihcob | 5´– GTT TTA CTG ATA GGT TAT TTA AAC-3´ | 1100 | Wicht et al. 2010 |
| R2Dnihcob | 5 – CAA TTT AAA AAA CGA GTT AAA GAT-3 | | |
| COBA | 5- GTA TGT GGC TGA TTC AGA GTT GAG C-3 | 400 | |
| COBB | 5-TTC GAG CCC GAA GAA TGC AAG TAG-3 | | Bouzid et al. 2008 |
| **COI** | | | |
| COIA2 | 5 -CAT ATG TTT TGA TTT TTT GG-3 | 400 | |
| COIB2 | 5-AKA ACA TAA TGA AAA TGA GC-3 | | Bouzid et al. 2008 |
| COIA2_L_Pavlovski | 5 -CAT ATG TTC TGG TTC TTT GG-3 | 400 | Present study |
| COIB2_L_Pavlovski | 5 -ATT ACA TAG TGA AAG TGA GC-3 | | |
| **ND1** | | | |
| Spi-ND1F | 5’ – GGA GAA TAT TGG TTT GTC TAA CCA-3 | 1000 | Eom et al. 2018 |
| Spi-ND1R | 5 – CCT TCT TAA CGT TAA CAG CAT TAC GAT-3 | | |

Table S3 Results of BIC substitution model selection in PartitionFinder2 for different partitions of the mtDNA dataset.

| | BEAST | IQtree | MrBayes | | |
|---|---|---|---|---|---|
| | | | Substitution model | nst | rates |
| Cytb-codon1 | TRN+I+G+X | TRN+I+G | GTR+I+G | 6 | Invgamma |
| Cytb-codon2 | HKY+I+X | HKY+I | HKY+I | 2 | Propinv |
| Cytb-codon3 | GTR+G+X | GTR+G | GTR+G | 6 | Gamma |
| COI-codon1 | TRN+I+G+X | TRN+I+G | GTR+I+G | 6 | Invgamma |
| COI-codon2 | HKY+I+X | HKY+I | HKY+I | 2 | Propinv |
| COI-codon3 | GTR+G+X | GTR+G | GTR+G | 6 | Gamma |
| ND1-Codon1 | GTR+G+X | GTR+G | GTR+G | 6 | Gamma |

| | | | | | |
|---|---|---|---|---|---|
| ND1-codon2 | TRN+I+G+X | TRN+I+G | GTR+I+G | 6 | Invgamma |
| ND1-codon3 | HKY+I+X | HKY+I | HKY+I | 2 | Propinv |

Table S4. Maximum likelihood estimates for six biogeography scenarios indicate the DIVALIKE+ J showed the optimal range evolution (i.e. maximize likelihood). Abbreviations, DEC, dispersal–extinction–cladogenesis; ln LnL, log-likelihood; K, parameters; d, dispersal; l, extinction; j, cladogenesis per-event weights; AIC, Akaike information criterion, AIC-wt: Weighted Akaike information criterion.

| Data set | Model | LnL | numparams | d | e | j | AIC | AIC_wt |
|---|---|---|---|---|---|---|---|---|
| mtDNA | DEC | -15.81 | 2 | 1.00E-12 | 1.00E-12 | 0 | 35.63 | 0.069 |
| | DEC+J | -13.47 | 3 | 1.00E-12 | 1.00E-12 | 0.11 | 32.94 | 0.26 |
| | DIVALIKE | -17.52 | 2 | 0.027 | 1.00E-12 | 0 | 39.04 | 0.012 |
| | **DIVALIKE+J** | **-12.69** | **3** | **1.00E-12** | **1.00E-12** | **0.1** | **31.37** | **0.58** |
| | BAYAREALIKE | -24.68 | 2 | 0.035 | 0.36 | 0 | 53.36 | 9.70E-06 |
| | BAYAREALIKE+J | -14.65 | 3 | 1.00E-07 | 1.00E-07 | 0.13 | 35.3 | 0.081 |
| ddRAD | DEC | -20.28 | 2 | 0.018 | 0.0044 | 0 | 44.55 | 0.001 |
| | DEC+J | -13.35 | 3 | 1.00E-12 | 1.00E-12 | 0.088 | 32.71 | 0.37 |
| | DIVALIKE | -18.36 | 2 | 0.03 | 1.00E-12 | 0 | 40.73 | 0.0066 |
| | **DIVALIKE+J** | **-13.05** | **3** | **1.00E-12** | **1.00E-12** | **0.09** | **32.1** | **0.5** |
| | BAYAREALIKE | -24.12 | 2 | 0.039 | 0.34 | 0 | 52.24 | 2.10E-05 |
| | BAYAREALIKE+J | -14.39 | 3 | 1.00E-07 | 1.00E-07 | 0.099 | 34.78 | 0.13 |

Table S5. Host ancestral reconstruction . AIC and BIC for each model

| | mtDNA | | ddRAD | |
|---|---|---|---|---|
| | AIC | BIC | AIC | BIC |
| **ER** | **35.264** | **35.662** | **26.806** | **27.204** |
| SYM | 52.389 | 56.369 | 36.606 | 38.993 |
| ARD | 66.283 | 74.240 | 43.863 | 48.638 |

Table S6 Output of Dtrios command: results of D statistics tests for 11 lineages. Trios with Z > 3 are shown in the Table.

| P1 | P2 | P3 | Dstatistic | Z-score | p-value | f4-ratio | BBAA | ABBA | BABA |
|---|---|---|---|---|---|---|---|---|---|
| L. alternans | LineageA | L. pavlovskii | 0.492497 | 9.65808 | 0 | 0.05994 | 49.7675 | 44.6564 | 15.1848 |
| LineageA | LineageB | L. pavlovskii | 0.353104 | 8.08807 | 6.66E-16 | 0.051701 | 50.1847 | 45.8005 | 21.8964 |
| Australia | LineageB | LineageA | 0.326624 | 6.69574 | 2.15E-11 | 0.155122 | 132.357 | 25.9863 | 13.1903 |
| NewZealand | LineageB | LineageA | 0.321417 | 6.55221 | 5.67E-11 | 0.151427 | 131.57 | 25.5616 | 13.1266 |
| L. pavlovskii | LineageB | China | 0.385941 | 6.28436 | 3.29E-10 | 0.099393 | 67.3717 | 57.2107 | 25.3479 |
| China | NewZealand | L. pavlovskii | 0.379399 | 5.99123 | 2.08E-09 | 0.071975 | 66.2316 | 62.8912 | 28.2952 |
| China | Australia | L. pavlovskii | 0.374515 | 5.86748 | 4.42E-09 | 0.071631 | 66.4985 | 63.1818 | 28.7515 |
| LineageA | L. alternans | Canada | 0.443981 | 5.86069 | 4.61E-09 | 0.059504 | 83.4059 | 38.5717 | 14.8524 |
| Canada | L. alternans | EAR | 0.461102 | 5.67512 | 1.39E-08 | 0.05135 | 66.5512 | 55.097 | 20.3214 |
| Canada | L. alternans | Africa | 0.454264 | 5.54966 | 2.86E-08 | 0.053939 | 65.35 | 56.7708 | 21.3042 |
| EAR | LineageA | Canada | 0.399662 | 5.47819 | 4.30E-08 | 0.053472 | 67.0639 | 39.4172 | 16.9067 |
| China | LineageA | L. pavlovskii | 0.278626 | 5.33177 | 9.73E-08 | 0.037697 | 48.1374 | 41.576 | 23.4563 |
| LineageB | L. alternans | Canada | 0.424636 | 5.18445 | 2.17E-07 | 0.064532 | 70.1938 | 43.424 | 17.5375 |
| LineageB | LineageA | L. alternans | 0.345835 | 4.92105 | 8.61E-07 | 0.044535 | 58.3266 | 35.1494 | 17.0849 |
| Africa | LineageA | Canada | 0.364392 | 4.89599 | 9.78E-07 | 0.048537 | 67.813 | 38.0544 | 17.7278 |
| China | LineageA | Ethiopia | 0.345126 | 4.63539 | 3.56E-06 | 0.016629 | 210.466 | 21.4168 | 10.4268 |
| L. alternans | China | LineageB | 0.309172 | 4.4105 | 1.03E-05 | 0.147855 | 70.4412 | 56.5223 | 29.8259 |
| EAR | China | Canada | 0.360416 | 4.38773 | 1.15E-05 | 0.069736 | 60.55 | 55.4054 | 26.0482 |
| L. pavlovskii | L. alternans | Canada | 0.337482 | 4.36498 | 1.27E-05 | 0.06389 | 60.9341 | 50.6826 | 25.1055 |
| L. alternans | China | NewZealand | 0.333782 | 4.16317 | 3.14E-05 | 0.067358 | 72.6845 | 64.8393 | 32.3869 |
| L. alternans | China | Australia | 0.331862 | 4.12352 | 3.73E-05 | 0.070369 | 73.0595 | 65.025 | 32.6202 |
| China | LineageB | EAR | 0.323237 | 4.09997 | 4.13E-05 | 0.03395 | 88.0403 | 44.6698 | 22.8461 |
| EAR | LineageB | Canada | 0.319196 | 4.08641 | 4.38E-05 | 0.048324 | 78.0203 | 42.0382 | 21.6949 |
| China | NewZealand | EAR | 0.354538 | 3.96094 | 7.47E-05 | 0.043836 | 96.9184 | 53.8291 | 25.6505 |
| Africa | China | Canada | 0.334574 | 3.9394 | 8.17E-05 | 0.064886 | 60.3399 | 54.1952 | 27.022 |
| China | LineageB | Africa | 0.312282 | 3.86498 | 0.000111 | 0.03337 | 87.1603 | 43.8233 | 22.9661 |
| China | LineageA | Africa | 0.294843 | 3.86012 | 0.000113 | 0.026913 | 77.0632 | 36.8179 | 20.0506 |
| Australia | L. alternans | Canada | 0.379397 | 3.81367 | 0.000137 | 0.062719 | 72.5625 | 45.6167 | 20.5233 |
| China | LineageA | EAR | 0.273006 | 3.80504 | 0.000142 | 0.024392 | 78.8571 | 36.5 | 20.8446 |
| L. pavlovskii | NewZealand | LineageA | 0.21155 | 3.7911 | 0.00015 | 0.162041 | 46.019 | 45.396 | 29.5427 |
| China | Australia | EAR | 0.336092 | 3.75798 | 0.000171 | 0.041517 | 97.3969 | 52.9648 | 26.3184 |
| EAR | L. pavlovskii | Canada | 0.289487 | 3.74482 | 0.000181 | 0.049059 | 79.3299 | 45.9975 | 25.3449 |
| L. pavlovskii | Australia | LineageA | 0.205122 | 3.71137 | 0.000206 | 0.158352 | 46.3283 | 45.5099 | 30.0176 |
| NewZealand | L. alternans | Canada | 0.375746 | 3.67872 | 0.000234 | 0.061676 | 72.2292 | 45.1417 | 20.4833 |
| China | NewZealand | Africa | 0.335035 | 3.64698 | 0.000265 | 0.042571 | 96.631 | 52.9286 | 26.3631 |
| L. alternans | LineageA | Ethiopia | 0.274298 | 3.5445 | 0.000393 | 0.011789 | 205.605 | 18.0065 | 10.2546 |
| Australia | LineageA | L. alternans | 0.279811 | 3.54036 | 0.0004 | 0.044203 | 51.2906 | 40.9094 | 23.021 |
| NewZealand | LineageA | L. alternans | 0.28029 | 3.51274 | 0.000444 | 0.043464 | 51.2734 | 40.5312 | 22.7845 |
| Africa | LineageB | Canada | 0.282448 | 3.51161 | 0.000445 | 0.043362 | 77.2417 | 41.226 | 23.0667 |
| China | Australia | Africa | 0.317076 | 3.49414 | 0.000476 | 0.040299 | 97.1167 | 52.1452 | 27.0381 |
| EAR | NewZealand | Canada | 0.324139 | 3.47698 | 0.000507 | 0.051242 | 85.1696 | 44.0607 | 22.4893 |
| L. pavlovskii | LineageB | L. alternans | 0.261776 | 3.45404 | 0.000552 | 0.039019 | 76.6668 | 39.8094 | 23.2912 |
| EAR | Australia | Canada | 0.313295 | 3.44283 | 0.000576 | 0.050208 | 84.3125 | 44.3007 | 23.1643 |
| L. alternans | LineageB | EAR | 0.27597 | 3.4073 | 0.000656 | 0.033589 | 66.4929 | 49.8188 | 28.269 |
| L. alternans | NewZealand | EAR | 0.3165 | 3.34234 | 0.000831 | 0.043427 | 68.6726 | 58.0357 | 30.131 |
| Africa | L. pavlovskii | Canada | 0.264706 | 3.33406 | 0.000856 | 0.044101 | 78.3521 | 44.1198 | 25.651 |
| L. alternans | LineageA | EAR | 0.244146 | 3.31422 | 0.000919 | 0.023975 | 80.0984 | 39.1916 | 23.81 |

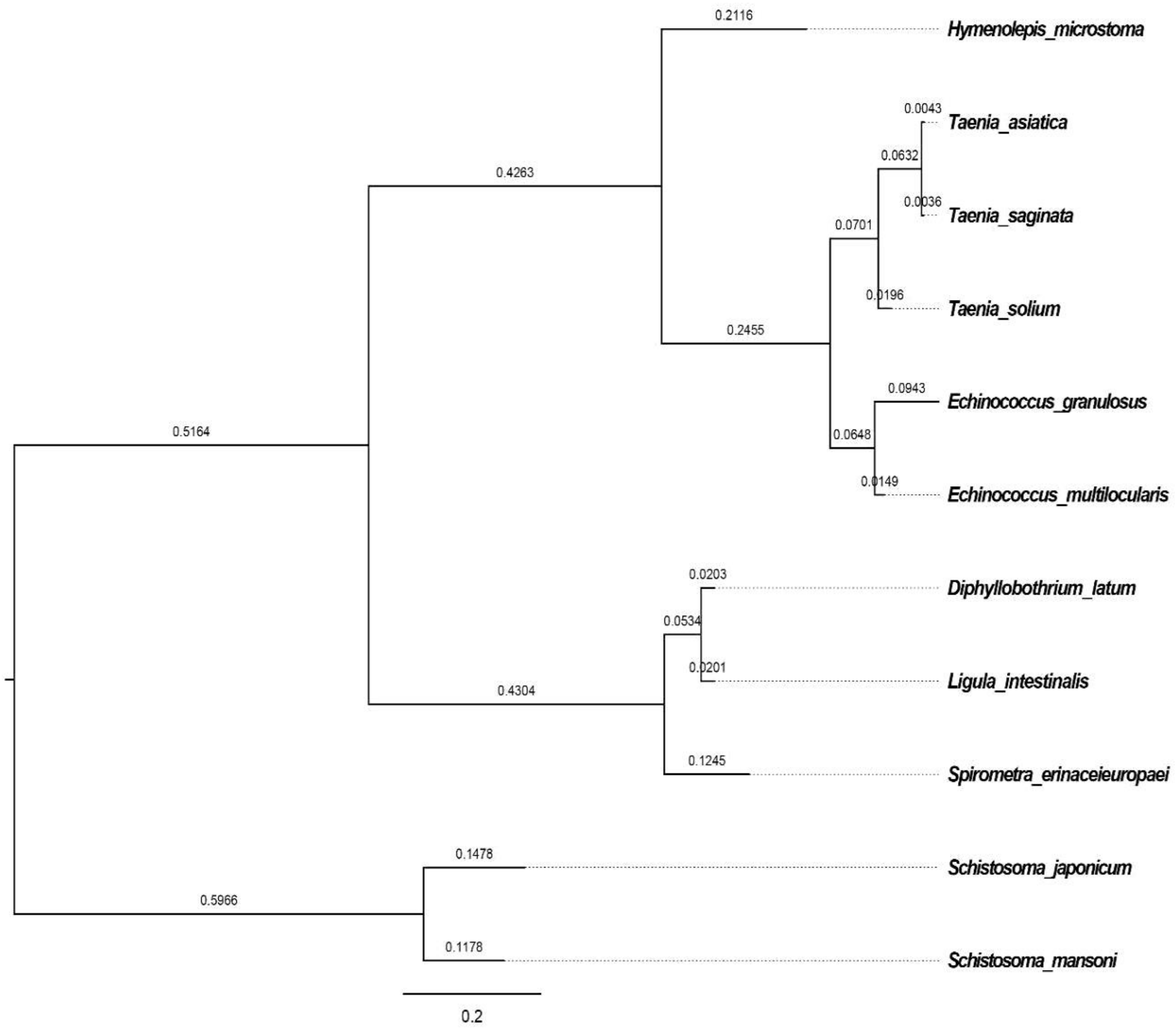


Fig. S1 phylogeny of 11 species using maximum likelihood inferred from the coding DNA sequences (CDS). Substitution rates are shown above the branches.

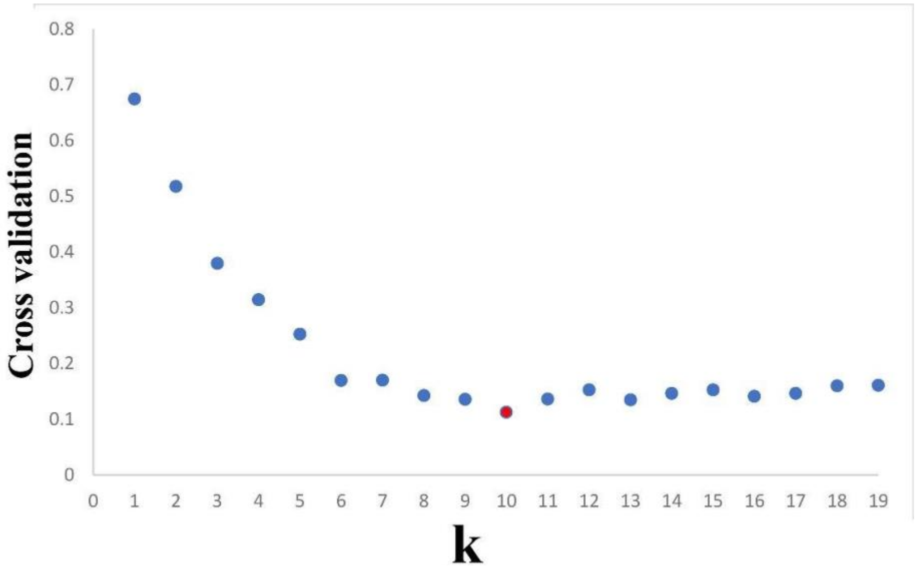


Fig. S2 Admixture cross validation from K=1 to K-19. K=10 was identified as the best predictive accuracy.

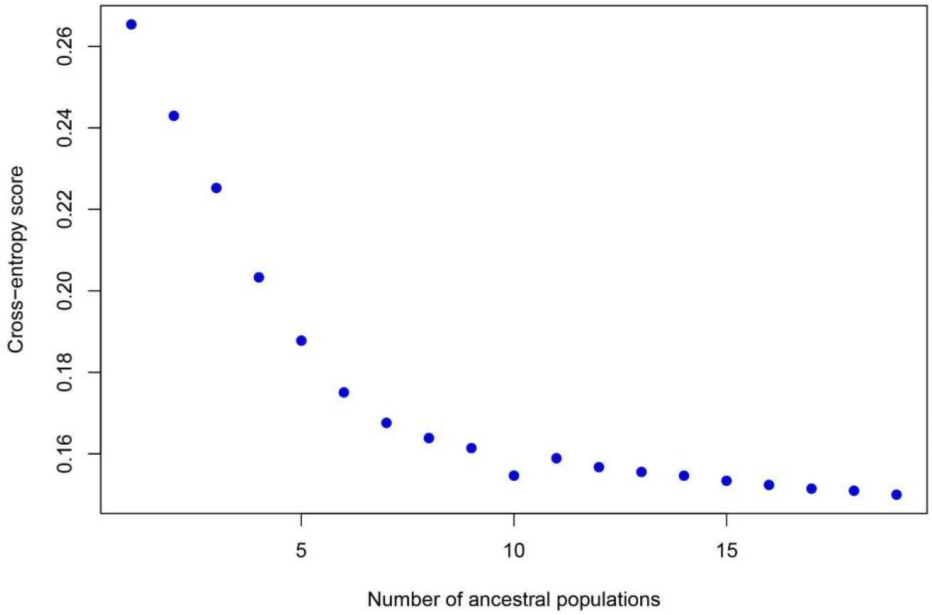


Fig. S3 Cross-entropy plot from the LEA r package revealed 10 genetic clusters for the *Ligula* complex species.

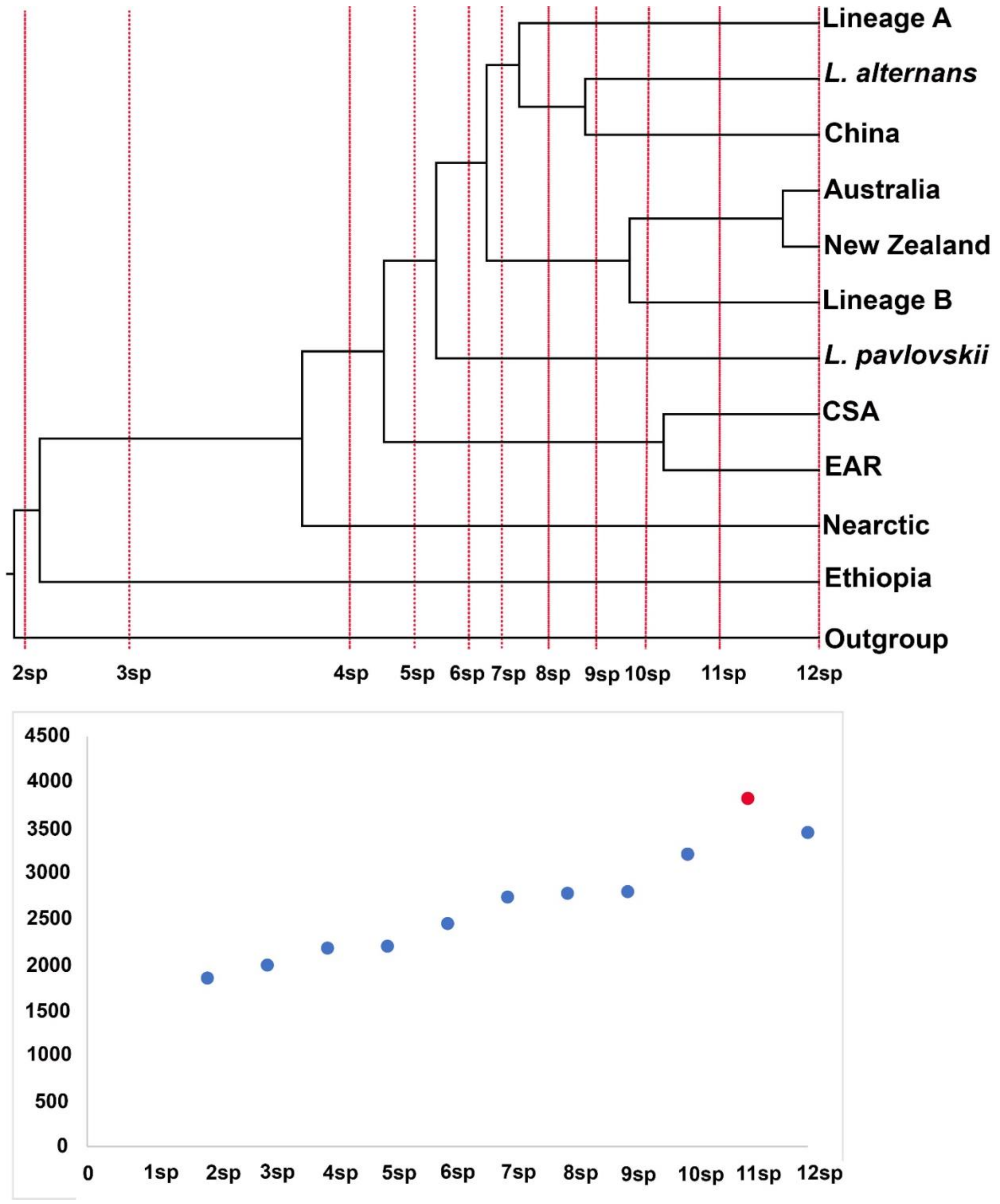


Fig. S4 Marginal-likelihood results for 12 species hypotheses based on the Bayes Factor Delimitation analysis.

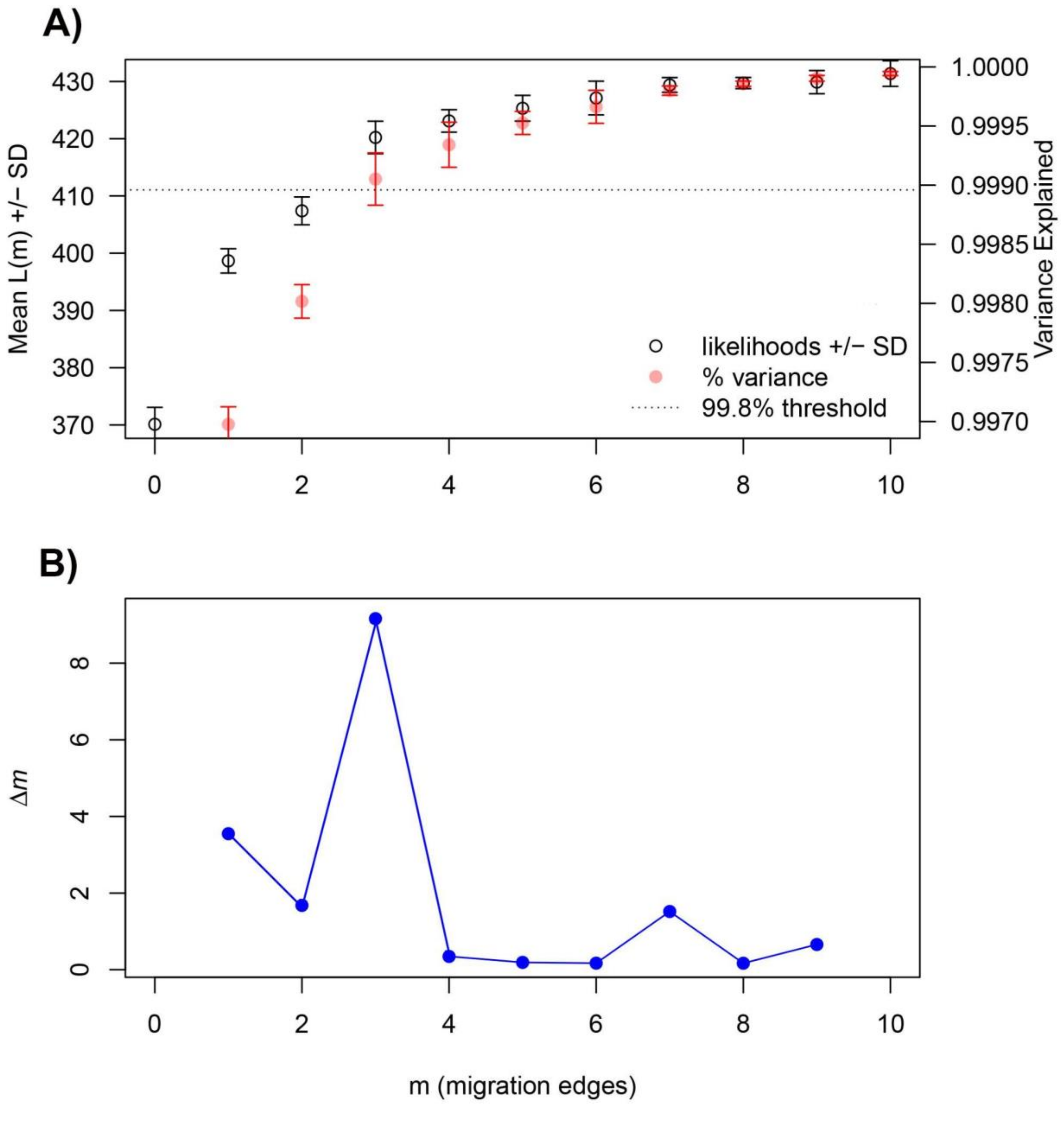


Fig. S5 Results of OptM reveal that the simplest model in TreeMix comprise three migrations routs. A) the black dotted line shows 99.8% variance explained by m=3 (it is recommended by Pickrell and Pritchard, 2012. B) The second-order rate of change (Dm) demonstrated the maximum value (9.3208) for delta at m = 3 edges.
