## Supplementary material for "Historical dispersal and host-switching formed the evolutionary history of a globally distributed multi-host parasite - the *Ligula intestinalis* species complex": Table S1a

| id | host | locality | Country | number of reads | Barcode | index | Accession number |
|---|---|---|---|---|---|---|---|
| Li3_L1 | *Blicca bjoerkna* | Lipno reservoir | Czech republic | 5486672 | CGATC | ATCACG | XXXXXXX |
| Li5_L1 | *Blicca bjoerkna* | Lipno reservoir | Czech republic | 5576052 | TCGAT | ATCACG | XXXXXXX |
| Li9_L1 | *Abramis brama* | Lipno reservoir | Czech republic | 5571409 | TGCAT | ATCACG | XXXXXXX |
| Li12_L1 | *Abramis brama* | Lipno reservoir | Czech republic | 4818508 | GGTTG | ATCACG | XXXXXXX |
| S2_L1 | *Esox lucius* | Medard reservoir | Czech republic | 6082925 | CGAAT | ATCACG | XXXXXXX |
| CF_L1 | *Blicca bjoerkna* | Medard reservoir | Czech republic | 6221424 | CGGCT | ATCACG | XXXXXXX |
| PA1_L1 | *Rutilus rutilus* | Medard reservoir | Czech republic | 4733833 | CGGTA | ATCACG | XXXXXXX |
| PE1_L1 | *Rutilus rutilus* | Medard reservoir | Czech republic | 6591323 | CGTAC | ATCACG | XXXXXXX |
| Af2_L1 | Engraulicypris sardella | Lake Malawi/Nyasa | Tanzania | 4468390 | CGTCG | ATCACG | XXXXXXX |
| Af4_L1 | Engraulicypris sardella | Lake Malawi/Nyasa | Tanzania | 4489443 | CTGAT | ATCACG | XXXXXXX |
| ATS77_L2 | Barbus humilis | Tana Lake, Bahir Dar | Ethiopia | 12020491 | GAGTC | ATCACG | XXXXXXX |
| ATS80_L2 | Barbus humilis | Tana Lake, Bahir Dar | Ethiopia | 13884205 | GCCGT | ATCACG | XXXXXXX |
| ATS84a_L2 | Barbus tsanensis | Tana Lake, Bahir Dar | Ethiopia | 13128679 | GCTGA | ATCACG | XXXXXXX |
| AU2_L2 | Galaxias maculatus | Goodga River, Western Australia | Austrialia | 11289209 | GGATA | ATCACG | XXXXXXX |
| AU3_L2 | Galaxias maculatus | Moates Lake, Western Australia | Austrialia | 16639212 | GGCCA | ATCACG | XXXXXXX |
| AU4_L2 | Galaxias maculatus | Moates Lake, Western Australia | Austrialia | 14310464 | GGCTC | ATCACG | XXXXXXX |
| AU5_L2 | Galaxias maculatus | Moates Lake, Western Australia | Austrialia | 14870785 | GTAGT | ATCACG | XXXXXXX |
| HU6_L2 | Neogobius fluviatilis | Balaton lake | Hungary | 13449952 | GTCCG | ATCACG | XXXXXXX |
| HU9_L2 | Neogobius fluviatilis | Balaton lake | Hungary | 16996354 | GTCGA | ATCACG | XXXXXXX |
| HU13_L2 | Neogobius fluviatilis | Balaton lake | Hungary | 14497443 | TACGT | ATCACG | XXXXXXX |
| Ba151_L2 | Barbus anoplus | Buffelskloof Spruit river | South Africa | 15006962 | TAGTA | ATCACG | XXXXXXX |
| DRC316_L2 | Barbus | Mwadingusha Lake | Democratic Republic of the Cong | 14473172 | TCAGT | ATCACG | XXXXXXX |
| NaH161A_L2 | Barbus paludinosus | Hardap dam | South Africa | 14527049 | TCCGG | ATCACG | XXXXXXX |
| NaH637_L2 | Barbus paludinosus | Hardap dam | South Africa | 15586050 | TCTGC | ATCACG | XXXXXXX |
| E82A_L3 | Barbus tsanensis | Tana Lake, Bahir Dar | Ethiopia | 8774995 | ACTTC | CGATGT | XXXXXXX |
| E82B_L3 | Barbus tsanensis | Tana Lake, Bahir Dar | Ethiopia | 6889676 | ATACG | CGATGT | XXXXXXX |
| E83A_L3 | Barbus tsanensis | Tana Lake, Bahir Dar | Ethiopia | 8040507 | ATGAG | CGATGT | XXXXXXX |
| E90_L3 | Barbus brevicephalus | Tana Lake, Bahir Dar | Ethiopia | 8081375 | ATTAC | CGATGT | XXXXXXX |
| E88A_L3 | Barbus intermedius | Tana Lake, Bahir Dar | Ethiopia | 6861512 | CATAT | CGATGT | XXXXXXX |
| AU8_L3 | Galaxias truttaceus | Moates Lake, Western Australia | Austrialia | 8609554 | CGGTA | CGATGT | XXXXXXX |
| CE2_L3 | Blicca bjoerkna | Medard reservoir | Czech republic | 10733223 | TCACG | CGATGT | XXXXXXX |
| PK_L3 | Rutilus rutilus | Medard reservoir | Czech republic | 9863929 | TCCGG | CGATGT | XXXXXXX |
| FR75_L4 | Rutilus rutilus | Créteil | France | 7353833 | GCATG | TTAGGC | XXXXXXX |
| PL8_L4 | Rhodeus amarus | Wloclawski Reservoir, Vistula | Poland | 8240713 | AACCA | TTAGGC | XXXXXXX |
| GB3_L4 | Phoxinus phoxinus | Aberystwyth, wales | United Kingdom | 7053097 | CGATC | TTAGGC | XXXXXXX |
| IE1_L4 | Rutilus rutilus | Lough Neagh | Ireland | 5816625 | TCGAT | TTAGGC | XXXXXXX |
| IE2_L4 | Rutilus rutilus | Lough Neagh | Ireland | 5905188 | TGCAT | TTAGGC | XXXXXXX |
| IE3_L4 | Gobio Gobio | Lough Neagh | Ireland | 5734870 | CAACC | TTAGGC | XXXXXXX |
| IE4_L4 | Gobio Gobio | Lough Neagh | Ireland | 5921707 | GGTTG | TTAGGC | XXXXXXX |
| RU1_L4 | Hemiculter lucidus | Lac Khanka | Russia | 7050501 | AAGGA | TTAGGC | XXXXXXX |
| CN1_L4 | Hemiculter bleekeri blee | lac Dong ting | China | 6753195 | AGCTA | TTAGGC | XXXXXXX |
| CN7_L4 | Neosalanx taihuensis | Tian' e Zhou (Zhanghe reserve | China | 8632402 | ACACA | TTAGGC | XXXXXXX |
| CN8_L4 | Neosalanx taihuensis | Tian' e Zhou (Zhanghe reserve | China | 5764805 | AATTA | TTAGGC | XXXXXXX |
| TANZ2_L4 | Engraulicypris sardella | Lake Malawi/Nyasa | Tanzania | 5456877 | ACGGT | TTAGGC | XXXXXXX |
| TANZ3_L4 | Engraulicypris sardella | Lake Malawi/Nyasa | Tanzania | 7270062 | ACTGG | TTAGGC | XXXXXXX |
| TANZ4_L4 | Engraulicypris sardella | Lake Malawi/Nyasa | Tanzania | 7088457 | ACTTC | TTAGGC | XXXXXXX |
| TANZ7_L4 | Engraulicypris sardella | Lake Malawi/Nyasa | Tanzania | 7261810 | ATACG | TTAGGC | XXXXXXX |
| TANZ8a_L4 | Engraulicypris sardella | Lake Malawi/Nyasa | Tanzania | 7241836 | ATGAG | TTAGGC | XXXXXXX |
| E2b_L4 | Abramis brama | Peipsi | Estonia | 5157082 | CATAT | TTAGGC | XXXXXXX |
| TR3_L4 | Rhodeus sericeus | Masukiye stream, Sapanca Lake | Turkey | 3260022 | CGAAT | TTAGGC | XXXXXXX |
| TS04.126f_L4 | Abramis brama | Rybinsk | Russia | 4826446 | CGGCT | TTAGGC | XXXXXXX |
| TS04.126g_L4 | Abramis brama | Rybinsk | Russia | 3160095 | CGGTA | TTAGGC | XXXXXXX |
| TS04.126h_L4 | Abramis brama | Rybinsk | Russia | 6194506 | CGTAC | TTAGGC | XXXXXXX |
| TS04.126a_L4 | Abramis brama | Rybinsk | Russia | 4163635 | CGTCG | TTAGGC | XXXXXXX |
| TS04.126b_L4 | Abramis brama | Rybinsk | Russia | 3546454 | CTGAT | TTAGGC | XXXXXXX |
| TS04.126c_L4 | Abramis brama | Rybinsk | Russia | 5453257 | CTGCG | TTAGGC | XXXXXXX |
| CN2_L4 | Hemiculter bleekeri blee | lac Dong ting | China | 6235103 | CTGTC | TTAGGC | XXXXXXX |
| CN22_L4 | Neosalanx taihuensis | Zhanghe reserve | China | 7971377 | CTTGG | TTAGGC | XXXXXXX |
| CN23_L4 | Neosalanx taihuensis | Zhanghe reserve | China | 8583495 | GACAC | TTAGGC | XXXXXXX |
| CN30_L4 | Neosalanx taihuensis | Zhanghe reserve | China | 8481191 | GAGAT | TTAGGC | XXXXXXX |
| CN32_L4 | Neosalanx taihuensis | Zhanghe reserve | China | 7580750 | GCCGT | TTAGGC | XXXXXXX |
| CA27_L4 | Semotilus atromaculatu | Lac Dumbo | Canada | 7144289 | GCTGA | TTAGGC | XXXXXXX |
| CA28_L4 | Semotilus atromaculatu | Lac Dumbo | Canada | 7735787 | GGATA | TTAGGC | XXXXXXX |
| CA30_L4 | Semotilus atromaculatu | Lac Dumbo | Canada | 6442109 | GGCCA | TTAGGC | XXXXXXX |
| CA31_L4 | Semotilus atromaculatu | Lac Dumbo | Canada | 8804902 | GGCTC | TTAGGC | XXXXXXX |
| UA1_L4 | Alburnus alburnus | Dniester | Ukraine | 6421321 | GTAGT | TTAGGC | XXXXXXX |
| UA2_L4 | Carassius carassius | Dniester | Ukraine | 5673661 | GTCCG | TTAGGC | XXXXXXX |
| UA3_L4 | Rutilus rutilus | Dniester | Ukraine | 7927417 | GTCGA | TTAGGC | XXXXXXX |
| CR22_L4 | Rutilus rutilus | Créteil | France | 7761401 | TACCG | TTAGGC | XXXXXXX |
| CR23_L4 | Rutilus rutilus | Créteil | France | 6568657 | TACGT | TTAGGC | XXXXXXX |
| CR24_L4 | Rutilus rutilus | Créteil | France | 6807948 | TAGTA | TTAGGC | XXXXXXX |
| CR25_L4 | Rutilus rutilus | Créteil | France | 6941629 | TATAC | TTAGGC | XXXXXXX |
| CR26_L4 | Rutilus rutilus | Créteil | France | 6179618 | TCACG | TTAGGC | XXXXXXX |
| X20M48 | Rutilus rutilus | Most | Czech Republic | 6035246 | CATAT | CGATGT | XXXXXXX |
| CAN11 | Semotilus atromaculatu | Lac Dumbo | Canada | 6234219 | CGAAT | CGATGT | XXXXXXX |
| CAN12 | Semotilus atromaculatu | Lac Dumbo | Canada | 6032387 | CGGCT | CGATGT | XXXXXXX |
| LIPC1 | Abramis brama | Lipno | Czech Republic | 6191387 | CGGTA | CGATGT | XXXXXXX |
| BaBsNov156 | Barbus anoplus | Buffelskloof Spruit river | South Africa | 6375377 | CGTAC | CGATGT | XXXXXXX |
| C6 | Rhinichthys osculus | Mckenzie River | Canada | 8112604 | CGTCG | CGATGT | XXXXXXX |
| X20M57 | Scardinius erythropthalr | Most | Czech Republic | 6310454 | CTGAT | CGATGT | XXXXXXX |
| S5_2 | Rutilus rutilus | Alborz Dam | Iran | 7117135 | CTGTC | CGATGT | XXXXXXX |
| S6_1 | Rutilus rutilus | Alborz Dam | Iran | 7005135 | CTTGG | CGATGT | XXXXXXX |
| S7_2 | Rutilus rutilus | Alborz Dam | Iran | 6202440 | GACAC | CGATGT | XXXXXXX |
| YK1 | Apollonia fluviatilis | Lake Kitay, near Danube River | Ukraine | 3365552 | GAGTC | CGATGT | XXXXXXX |
| YK2 | Apollonia fluviatilis | Lake Kitay, near Danube River | Ukraine | 5132802 | GCCGT | CGATGT | XXXXXXX |
| YK26 | Pomatoschistus microps | Unterwarnow, near Kleine Warnow | Germany | 4801776 | GCTGA | CGATGT | XXXXXXX |
| YK119 | Pomatoschistus microps | Gdynia, Gulf of Gdańsk | Poland | 6802708 | GGATA | CGATGT | XXXXXXX |
| YK131 | Apollonia fluviatilis | Dniester River | Ukraine | 6953413 | GGCCA | CGATGT | XXXXXXX |
| YK297 | Neogobius melanostomu | Rvach River | Ukraine | 6647697 | GGCTC | CGATGT | XXXXXXX |
| YK308 | Neogobius fluviatilis | Sasyk Lagoon | Ukraine | 6640430 | GTAGT | CGATGT | XXXXXXX |
| POL5 | Rhodeus sericeus | Wlodawski reservoir | Poland | 6490951 | GTCCG | CGATGT | XXXXXXX |
| Bal3 | Neogobius fluviatilis | Balaton lake | Hungary | 6621613 | TACCG | CGATGT | XXXXXXX |
| Bal4 | Neogobius fluviatilis | Balaton lake | Hungary | 5649532 | TACGT | CGATGT | XXXXXXX |
| Bal6 | Neogobius fluviatilis | Balaton lake | Hungary | 5605614 | TAGTA | CGATGT | XXXXXXX |
| Bal7 | Neogobius fluviatilis | Balaton lake | Hungary | 5764920 | TATAC | CGATGT | XXXXXXX |
| FR77 | Rutilus rutilus | Créteil | France | 4532929 | GCATG | ATCACG | XXXXXXX |
| FR88 | Rutilus rutilus | Créteil | France | 6005983 | AACCA | ATCACG | XXXXXXX |
| MPL28 | Rutilus rutilus | Most | Czech Republic | 6509615 | CGATC | ATCACG | XXXXXXX |
| FR84 | Rutilus rutilus | Créteil | France | 5568505 | TCGAT | ATCACG | XXXXXXX |
| FR89 | Rutilus rutilus | Créteil | France | 4560249 | TGCAT | ATCACG | XXXXXXX |
| MPL29 | Rutilus rutilus | Most | Czech Republic | 5019649 | CAACC | ATCACG | XXXXXXX |
| MPL30 | Rutilus rutilus | Most | Czech Republic | 4529673 | GGTTG | ATCACG | XXXXXXX |
| X2Ab | Abramis brama | Římov | Czech Republic | 3418401 | AAGGA | ATCACG | XXXXXXX |
| X3Ab | Abramis brama | Římov | Czech Republic | 4630225 | AGCTA | ATCACG | XXXXXXX |
| X4Ab | Abramis brama | Římov | Czech Republic | 3987628 | ACACA | ATCACG | XXXXXXX |
| X9Rr | Rutilus rutilus | Římov | Czech Republic | 4100447 | AATTA | ATCACG | XXXXXXX |
| X16Ab | Abramis brama | Římov | Czech Republic | 3375762 | ACGGT | ATCACG | XXXXXXX |
| TR7 | Rhodeus sericeus | Kurtköy Stream, Sapanca Lake | Turkey | 5811174 | ACTGG | ATCACG | XXXXXXX |
| TR8 | Rhodeus sericeus | Kurtköy Stream, Sapanca Lake | Turkey | 5638767 | ACTTC | ATCACG | XXXXXXX |
| CA7 | Rhodeus sericeus | Lac Dumbo | Canada | 5093620 | ATACG | ATCACG | XXXXXXX |
| TS76 | Barbus humilis | Tana Lake, Bahir Dar | Ethiopia | 4225532 | ATGAG | ATCACG | XXXXXXX |
| TS79 | Barbus tsanensis | Tana Lake, Bahir Dar | Ethiopia | 4090140 | ATTAC | ATCACG | XXXXXXX |
| TS81a | Barbus intermedius | Tana Lake, Bahir Dar | Ethiopia | 4308927 | CATAT | ATCACG | XXXXXXX |
| C1 | Rhinichthys osculus | Mckenzie River | Canada | 3118397 | CGAAT | ATCACG | XXXXXXX |
| C3 | Rhinichthys osculus | Mckenzie River | Canada | 5345730 | CGGCT | ATCACG | XXXXXXX |
| Li2 | Blicca bjoerkna | Lipno | Czech Republic | 5036084 | CGGTA | ATCACG | XXXXXXX |
| Li14 | Abramis brama | Lipno | Czech Republic | 5286573 | CGTAC | ATCACG | XXXXXXX |
| X20R1PL | Rutilus rutilus | Římov | Czech Republic | 5850754 | CGTCG | ATCACG | XXXXXXX |
| AB1 | Blicca bjoerkna | Římov | Czech Republic | 5028862 | CTGAT | ATCACG | XXXXXXX |
| AB2 | Blicca bjoerkna | Římov | Czech Republic | 4445081 | CTGCG | ATCACG | XXXXXXX |
| AbLipno1 | Abramis brama | Lipno | Czech Republic | 5418712 | GAGAT | ATCACG | XXXXXXX |
| AbLipno2 | Abramis brama | Lipno | Czech Republic | 5094581 | GAGTC | ATCACG | XXXXXXX |
| X20M11 | Rutilus rutilus | Most | Czech Republic | 5084222 | GCCGT | ATCACG | XXXXXXX |
| X20M34 | Rutilus rutilus | Most | Czech Republic | 5876404 | GCTGA | ATCACG | XXXXXXX |
| X20M51 | Rutilus rutilus | Most | Czech Republic | 4112140 | GTAGT | ATCACG | XXXXXXX |
| X20M58 | Scardinius erythropthalr | Most | Czech Republic | 5502583 | GTCCG | ATCACG | XXXXXXX |
| Hu8 | Neogobius fluviatilis | Balaton lake | Hungry | 5692982 | GTCGA | ATCACG | XXXXXXX |
| Hu10 | Neogobius fluviatilis | Balaton lake | Hungry | 5850812 | TACCG | ATCACG | XXXXXXX |
| Ea2 | Abramis brama | Peipsi | Estonia | 4480806 | TAGTA | ATCACG | XXXXXXX |
| NZ2 | Gobiomorphus breviceps | | New Zealand | 5434883 | TCACG | ATCACG | XXXXXXX |
| MPR1 | Scardinius erythrophtha | Most | Czech Republic | 6283789 | TCAGT | ATCACG | XXXXXXX |
| MPR2 | Scardinius erythrophtha | Most | Czech Republic | 5779666 | TCCGG | ATCACG | XXXXXXX |
| MPR3 | Scardinius erythrophtha | Most | Czech Republic | 5855323 | TCTGC | ATCACG | XXXXXXX |
| CN31 | Neosalanx taihuensis | Zhanghe reserve | China | 6659308 | TGGAA | ATCACG | XXXXXXX |
| MPL27B | Rutilus rutilus | Most | Czech Republic | 5807870 | TTACC | ATCACG | XXXXXXX |
| E88b_L4 | Barbus intermedius | Tana Lake, Bahir Dar | Ethiopia | 6659308 | TCTGC | ATCACG | XXXXXXX |
| Ba159_L3 | Barbus anoplus | Buffelskloof Spruit river | South Africa | 10165369 | CGTAC | CGATGT | XXXXXXX |
| TU61_L3 | Pseudophoxinus callensi | Joumine, Ichkeul | Tunisia | 8293854 | CGTCG | CGATGT | XXXXXXX |
| Ir8_L3 | Squalius orientalis | | Iran | 8054152 | GCTGA | CGATGT | XXXXXXX |
| TOLAR | | | | 3421276 | TGGAA | CGATGT | XXXXXXX |
| CHIL2 | | | | 1521821 | TTACC | CGATGT | XXXXXXX |
