## Supplementary material for "Historical dispersal and host-switching formed the evolutionary history of a globally distributed multi-host parasite - the *Ligula intestinalis* species complex": Table S1b

| id | Cytb | COI | ND1 | Host | | localiy | refernce |
|---|---|---|---|---|---|---|---|
| Au1 | xxxxxx | xxxxxx | xxxxxx | *Galaxias maculatus* | Australia | Goodga Ri | Present Study |
| Au2 | xxxxxx | xxxxxx | xxxxxx | *Galaxias maculatus* | Australia | Goodga Ri | Present Study |
| Au3 | xxxxxx | xxxxxx | xxxxxx | *Galaxias maculatus* | Australia | Moates Lal | Present Study |
| Au5 | xxxxxx | xxxxxx | xxxxxx | *Galaxias maculatus* | Australia | Moates Lal | Present Study |
| Au6 | xxxxxx | xxxxxx | xxxxxx | *Galaxias maculatus* | Australia | Moates Lal | Present Study |
| Au7 | xxxxxx | xxxxxx | xxxxxx | *Galaxias maculatus* | Australia | Moates Lal | Present Study |
| Au8 | xxxxxx | xxxxxx | xxxxxx | *Galaxias maculatus* | Australia | Moates Lal | Present Study |
| Au9 | xxxxxx | xxxxxx | xxxxxx | *Galaxias maculatus* | Australia | Moates Lal | Present Study |
| YK1 | xxxxxx | xxxxxx | xxxxxx | *Apollonia fluviatilis* | Ukraine | Lake Kitay | Present Study |
| YK2 | xxxxxx | xxxxxx | xxxxxx | *Apollonia fluviatilis* | Ukraine | Lake Kitay | Present Study |
| HU10 | xxxxxx | xxxxxx | xxxxxx | *Neogobius fluviatilis* | Hungary | Balaton lak | Present Study |
| Bal3 | xxxxxx | xxxxxx | xxxxxx | *Neogobius fluviatilis* | Hungary | Balaton lak | Present Study |
| Bal4 | xxxxxx | xxxxxx | xxxxxx | *Neogobius fluviatilis* | Hungary | Balaton lak | Present Study |
| yk30 | xxxxxx | xxxxxx | xxxxxx | *Neogobius fluviatilis* | Ukraine | Sasyk Lag | Present Study |
| yk69 | xxxxxx | xxxxxx | xxxxxx | *Neogobius fluviatilis* | Ukraine | Dnistrovsk | Present Study |
| YK119 | xxxxxx | xxxxxx | xxxxxx | *Pomatoschistus minutus* | Poland | Gdynia, G | Present Study |
| yk131 | xxxxxx | xxxxxx | xxxxxx | *Apollonia fluviatilis* | Ukraine | Dniester R | Present Study |
| yk202 | xxxxxx | xxxxxx | xxxxxx | *Pomatoschistus microps* | Germany | Unterwarn | Present Study |
| CZ121Mm | EU241193 | EU241239 | | *Mergus merganser* | Czech Republic | Bolevec, p | Bouzid et al., 2008 |
| 126a | xxxxxx | xxxxxx | xxxxxx | *Abramis brama* | Russia | Rybinsk | Present Study |
| 126c | xxxxxx | xxxxxx | xxxxxx | *Abramis brama* | Russia | Rybinsk | Present Study |
| 126g | xxxxxx | xxxxxx | xxxxxx | *Abramis brama* | Russia | Rybinsk | Present Study |
| 126h | xxxxxx | xxxxxx | xxxxxx | *Abramis brama* | Russia | Rybinsk | Present Study |
| S4 | xxxxxx | xxxxxx | xxxxxx | *Rutilus rutilus* | Iran | Alborz Da | Present Study |
| S5 | xxxxxx | xxxxxx | xxxxxx | *Rutilus rutilus* | Iran | Alborz Da | Present Study |
| S6 | xxxxxx | xxxxxx | xxxxxx | *Rutilus rutilus* | Iran | Alborz Da | Present Study |
| S7 | xxxxxx | xxxxxx | xxxxxx | *Rutilus rutilus* | Iran | Alborz Da | Present Study |
| Ir8 | xxxxxx | xxxxxx | xxxxxx | *Squalius orientalis* | Iran | Zarde Lime | Present Study |
| OK | xxxxxx | xxxxxx | xxxxxx | *Rutilus rutilus* | Czech Republic | Medard | Present Study |
| CR22 | xxxxxx | xxxxxx | xxxxxx | *Rutilus rutilus* | France | Créteil | Present Study |
| CR23 | xxxxxx | xxxxxx | xxxxxx | *Rutilus rutilus* | France | Créteil | Present Study |
| It5 | xxxxxx | xxxxxx | xxxxxx | *Rutilus aula* | Italy | lago magg | Present Study |
| UA1 | xxxxxx | xxxxxx | xxxxxx | *Alburnus alburnus* | Ukraine | Dniester | Present Study |
| UA2 | xxxxxx | xxxxxx | xxxxxx | *Carassius carassius* | Ukraine | Dniester | Present Study |
| UA3 | xxxxxx | xxxxxx | xxxxxx | *Rutilus rutilus* | Ukraine | Dniester | Present Study |
| CZ106Pc | EU241167 | EU241244 | | *Podiceps cristatus* | Czech Republic | Zahlinice, S | Bouzid et al., 2008 |
| CZ90Rr | EU241178 | EU241282 | | *Rutilus rutilus* | Czech Republic | Zelivka, Bo | Bouzid et al., 2008 |
| CZ122Rs | | | | *Rutilus rutilus* | Czech Republic | Bolevec, p | Bouzid et al., 2008 |
| CZ10Rr | EU241165 | EU241280 | | *Rutilus rutilus* | Czech Republic | Lipno,s.Bo | Bouzid et al., 2008 |
| CZ24Ab | EU241180 | EU241267 | | *Abramis brama* | Czech Republic | Mlyny,s.m | Bouzid et al., 2008 |
| CZ16Ab | EU241179 | EU241283 | | *Abramis brama* | Czech Republic | Mlyny,s.m | Bouzid et al., 2008 |
| CZ6Rr | EU241169 | EU241277 | | *Rutilus rutilus* | Czech Republic | Lipno,s.Bo | Bouzid et al., 2008 |
| CZ7Rr | EU241159 | EU241278 | | *Rutilus rutilus* | Czech Republic | Lipno,s.Bo | Bouzid et al., 2008 |
| CZ18Ab | EU241184 | EU241285 | | *Abramis brama* | Czech Republic | Mlyny,s.m | Bouzid et al., 2008 |
| DE4Rr | EU241202 | EU241273 | | *Rutilus rutilus* | Germany | Lac Müggel | Bouzid et al., 2008 |
| CZ114Pc | EU241187 | EU241287 | | *Podiceps cristatus* | Czech Republic | Zahlinice, S | Bouzid et al., 2008 |
| EE1Ab | JQ279121 | JQ279085 | | *Abramis brama* | Estonia | Peipsi | Bouzid et al., 2013 |
| DE2Rr | JQ279116 | JQ279080 | | *Rutilus rutilus* | Germany | Lac Müggel | Bouzid et al., 2013 |
| CZ69Aa | EU241177 | EU241246 | | *Alburnus alburnus* | Czech Republic | Zelivka, Bo | Bouzid et al., 2008 |
| DE1Rr | EU241203 | EU241301 | | *Rutilus rutilus* | Germany | Lac Müggel | Bouzid et al., 2008 |
| FR90Rr | EU241172 | EU241299 | | *Rutilus rutilus* | France | Créteil | Bouzid et al., 2008 |
| CZ45Pc | EU241186 | EU241288 | | *Podiceps cristatus* | czech Republic | Tovacov | Bouzid et al., 2008 |
| DE6Rr | JQ279120 | JQ279084 | | *Rutilus rutilus* | Germany | Lac Mügge | Bouzid et al., 2013 |
| EE5Ab | JQ279122 | JQ279086 | | *Abramis brama* | Estonia | Peipsi | Bouzid et al., 2013 |
| FR7Rr | JQ279125 | JQ279090 | | *Rutilus rutilus* | France | Pareloup | Bouzid et al., 2013 |
| EE3ab | EU241160 | EU241276 | | *Abramis brama* | Estonia | Peipsi | Bouzid et al., 2008 |
| FR67Aa | JQ279124 | JQ279088 | | *Alburnus alburnus* | France | Pareloup | Bouzid et al., 2013 |
| EE4Ab | EU241195 | EU241294 | | *Abramis brama* | Estonia | Peipsi | Bouzid et al., 2013 |
| DE3Rr | EU241204 | EU241302 | | *Rutilus rutilus* | Germany | Lac Mügge | Bouzid et al., 2013 |
| IE6Rr | EU241207 | EU241249 | | *Rutilus rutilus* | Ireland | Lough Neag | Bouzid et al., 2013 |
| FR89Rr | JQ279126 | EU240998 | | *Rutilus rutilus* | France | Créteil | Bouzid et al., 2008 |
| GB1Rr | EU241205 | EU241303 | | *Rutilus rutilus* | United Kingdom | Scotland | Bouzid et al., 2008 |
| FR30Ab | EU241201 | EU241259 | | *Abramis brama* | France | Lavernose | Bouzid et al., 2008 |
| TN26Rb | JQ279127 | JQ279091 | | *Rutilus rubilio* | Tunisia | Sidi Salem | Bouzid et al., 2013 |
| FR91Rr | EU241173 | EU241300 | | *Rutilus rutilus* | France | Créteil | Bouzid et al., 2008 |
| RU7Ab | EU241255 | EU241075 | | *Abramis brama* | Russia | Rybinsk | Bouzid et al., 2008 |
| FR11Rr | EU241199 | EU241258 | | *Rutilus rutilus* | France | Pareloup | Bouzid et al., 2008 |
| UA3Rr | EU241176 | EU241317 | | *Rutilus rutilus* | Ukraine | Dniester | Bouzid et al., 2008 |
| CZ12Rr | EU241281 | EU241024 | | *Rutilus rutilus* | Czech republic | Lipno,s.Bo | Bouzid et al., 2013 |
| CZ14Ab | EU241182 | EU241263 | | *Abramis brama* | Czech republic | Mlyny,s.m | Bouzid et al., 2013 |
| RU5Ab | EU241210 | EU241309 | | *Abramis brama* | Russia | Rybinsk | Bouzid et al., 2013 |
| FR66Aa | JQ279123 | JQ279087 | | *Alburnus alburnus* | France | Pareloup | Bouzid et al., 2008 |
| RU8Ab | EU241211 | EU241310 | | *Abramis brama* | Russia | Rybinsk | Bouzid et al., 2013 |
| CZ17Ab | EU241183 | EU241284 | | *Abramis brama* | Czech republic | Mlyny,s.m | Bouzid et al., 2013 |
| GB2Pp | EU241175 | EU241304 | | *Phoxinus phoxinus* | United kingdom | River Gryfe | Bouzid et al., 2013 |
| TN58Se | JQ279131 | JQ279094 | | *Scardinus erythrophthalmus* | Tunisia | Nebhana | Bouzid et al., 2013 |
| H3 | xxxxx | xxxxx | | *Rutilus rutilus* | Hungary | Balaton | present study |
| TN50Se | JQ279130 | JQ279093 | | *Scardinus erythrophthalmus* | Tunisia | Sidi Salem | Bouzid et al., 2013 |
| UA2Cc | EU241218 | EU241238 | | *Carassius carassius* | Ukraine | Dniester | Bouzid et al., 2008 |
| 126b | xxxxx | xxxxx | | *Abramis brama* | Russia | | present study |
| TN60Rb | JQ279136 | JQ279099 | | *Rutilus rubilio* | Tunisia | Nebhana | Bouzid et al., 2013 |
| TU40Rr | xxxxxxx | xxxxxx | | *Rutilus rubilio* | Tunisia | Sidi Salem reservoir | |
| TN59Rb | JQ279135 | JQ279098 | | *Rutilus rubilio* | Tunisia | Nebhana | Bouzid et al., 2013 |
| TN66Se | JQ279141 | JQ279103 | | *Scardinus erythrophthalmus* | Tunisia | Sidi Salem | Bouzid et al., 2013 |
| UA1Aa | EU241181 | EU241316 | | *Alburnus alburnus* | Ukraine | Dniester | Bouzid et al., 2013 |
| IE2Rr | EU241206 | EU241250 | | *Rutilus rutilus* | Ireland | Lough Neagh, N. Ireland | |
| RU1CY | EU241209 | EU241311 | | *Hemiculter lucidus* | Russia | Lac Khanka | |
| IR3 | XXXX | XXXX | | *Cyprinus carpio* | Iran | Chaghakho | present study |
| IR1 | XXXXX | XXXX | | *Cyprinus carpio* | Iran | Chaghakho | present study |
| IR1N | XXXXX | XXXXX | | *Cyprinus carpio* | Iran | Chaghakho | present study |
| CN2 | XXXXX | XXXXX | XXXXX | *Hemiculter bleekeri* | China | lac Dong ti | present study |
| CN1Hb | EU241153 | EU241229 | | *Hemiculter bleekeri* | China | lac Dong ting | |
| CN4Nt | EU241157 | EU241237 | | *Neosalanx taihuensis* | China | Tian' e Zhou (Zhanghe reserve) | |
| CN16Nt | EU241156 | EU241236 | | *Neosalanx taihuensis* | China | Tian' e Zhou (Zhanghe reserve) | |
| CZ104Pc | EU241191 | EU241291 | | *Podiceps cristatus* | Czech Republic | Zahlinice, S.moravie | |
| TR5 | XXXXX | XXXXX | | *Rhodeus sericeus* | Turkey | Kurtköy St | present study |
| H11 | XXXXX | XXXXX | | *Neogobious* | Balaton lake | | present study |
| ALG1Bc | JQ279109 | JQ279074 | | *Barbus sp* | Algeria | | |
| IE4Gg | EU241208 | EU241290 | | *Gobio Gobio* | Ireland | Lough Neagh, N. Ireland | |
| ALG10Bc | JQ279106 | JQ279068 | | *Barbus sp* | Algeria | oued Hami | present study |
| TN62Ps | JQ279138 | JQ279101 | | *Pseudophoxinus callensis* | Tunisia | Joumine, Ichkeul | |
| TN61Ps | JQ279137 | JQ279100 | | *Pseudophoxinus callensis* | Tunisia | Joumine, Ichkeul | |
| ALG2Bc | EU241143 | EU241219 | | *Barbus sp* | Algeria | oued Hami | present study |
| IE3Gg | EU241188 | EU241305 | | *Gobio gobio* | Ireland | Lough Neagh, N. Ireland | |
| TN63Ps | JQ279139 | JQ279102 | | *Pseudophoxinus callensis* | Tunisia | Remel, Hammamet | |
| CZ2Pc | EU241190 | EU241293 | | *Podiceps cristatus* | Czech Republic | Zahlinice, S.moravie | |
| FR76Rr | EU241188 | EU241305 | | *Rutilus rutilus* | France | Créteil | present study |
| ALG12Bs | JQ279108 | JQ279073 | | *Barbus setivimensis* | Algeria | | |
| ALG9Bc | EU241188 | EU241305 | | *Barbus sp* | Algeria | | |
| PL8 | xxx | xxxx | | *Rhodeus amarus* | Poland | Wloclawsk | present study |
| Ru | xxxx | xxxx | | *Hemiculter lucidus* | Russia | Lac Khanka | present study |
| TR4 | xxxxxx | xxxxxx | xxxxxx | *Rhodeus sericeus* | Turkey | Kurtköy St | present study |
| AU4Gm | EU241147 | EU241223 | | *Galaxias truttaceus* | Australia | Goodga River, Western Australia | |
| CN8 | xxxxxx | xxxxxx | xxxxxx | Neosalanx taihuensis | China | | present study |
| CN31 | xxxxxx | xxxxxx | xxxxxx | Neosalanx taihuensis | China | | present study |
| CN30 | xxxxxx | xxxxxx | xxxxxx | Neosalanx taihuensis | China | | present study |
| AU1Gt | EU241146 | EU241222 | | Galaxias truttaceus | Australia | | |
| ALG6Bc | EU241188 | EU241305 | | | Algeria | | |
| ET2Bh | EU241196 | EU241295 | | | Ethiopia | | |
| ET2 | xxxxxx | xxxxxx | xxxxxx | | Ethiopia | | present study |
| ET5 | xxxxxx | xxxxxx | xxxxxx | | Ethiopia | | present study |
| ET7 | xxxxxx | xxxxxx | xxxxxx | | Ethiopia | | present study |
| ET5Bt | EU241197 | EU241296 | | | Ethiopia | | present study |
| ET7Bi | EU241198 | EU241297 | | | Ethiopia | | present study |
| Oregon_C1 | xxxxxx | xxxxxx | xxxxxx | Rhinichthys osculus | Canada | Mckenzie I | present study |
| CA2Sa | EU241148 | EU241225 | | Semotilus atromaculatus | Canada | | |
| Oregon_C3 | xxxxxx | xxxxxx | xxxxxx | Rhinichthys osculus | Canada | Mckenzie I | present study |
| CA19Sa | EU241150 | EU241224 | | Semotilus atromaculatus | Canada | | present study |
| CA5Sa | EU241149 | EU241226 | | Semotilus atromaculatus | Canada | | present study |
| Oregon_C4 | xxxxxx | xxxxxx | xxxxxx | Rhinichthys osculus | Canada | Mckenzie I | present study |
| CA1Cp | EU241152 | EU241228 | | *Couesius plumbeus* | Canada | | present study |
| CA14Sa | EU241151 | EU241227 | | | Canada | | present study |
| CA1 | xxxxxx | xxxxxx | xxxxxx | Semotilus atromaculatus | Canada | Lac Dumb | present study |
| CA15 | xxxxxx | xxxxxx | xxxxxx | Semotilus atromaculatus | Canada | Lac Dumb | present study |
| CA19 | xxxxxx | xxxxxx | xxxxxx | Semotilus atromaculatus | Canada | Lac Dumb | present study |
| K1CO | xxxxxx | xxxxxx | xxxxxx | | Kenya | | present study |
| 1c | xxxxxx | xxxxxx | xxxxxx | Rastrineobola argentea | Kenya | Victoria lak | present study |
| C2 | xxxxxx | xxxxxx | xxxxxx | Rastrineobola argentea | Kenya | Victoria lak | present study |
| C3 | xxxxxx | xxxxxx | xxxxxx | Rastrineobola argentea | Kenya | Victoria lak | present study |
| 4c | xxxxxx | xxxxxx | xxxxxx | Rastrineobola argentea | Kenya | Victoria lak | present study |
| 5c | xxxxxx | xxxxxx | xxxxxx | Rastrineobola argentea | Kenya | Victoria lak | present study |
| 6c | xxxxxx | xxxxxx | xxxxxx | Rastrineobola argentea | Kenya | Victoria lak | present study |
| 7c | xxxxxx | xxxxxx | xxxxxx | Rastrineobola argentea | Kenya | Victoria lak | present study |
| 8c | xxxxxx | xxxxxx | xxxxxx | Rastrineobola argentea | Kenya | Victoria lak | present study |
| 9c | xxxxxx | xxxxxx | xxxxxx | Rastrineobola argentea | Kenya | Victoria lak | present study |
| 10c | xxxxxx | xxxxxx | xxxxxx | Rastrineobola argentea | Kenya | Victoria lak | present study |
| K3CO | xxxxxx | xxxxxx | xxxxxx | | Kenya | | present study |
| K4CO | xxxxxx | xxxxxx | xxxxxx | | Kenya | | present study |
| Tanz2 | xxxxxx | xxxxxx | xxxxxx | Engraulicypris sardella | Tanzania | Lake Malawi/Nyasa | |
| Tanz3 | xxxxxx | xxxxxx | xxxxxx | Engraulicypris sardella | Tanzania | Lake Malawi/Nyasa | |
| Tanz8a | xxxxxx | xxxxxx | xxxxxx | Engraulicypris sardella | Tanzania | Lake Malawi/Nyasa | |
| Bn151 | xxxxxx | xxxxxx | xxxxxx | *Barbus anoplus* | South Africa | Buffelskloof Spruit river | |
| SA_Mpum | xxxxxx | xxxxxx | xxxxxx | | | | |
| DRC | xxxxxx | xxxxxx | xxxxxx | Barbus sp | Democratic Republic | Mwadingusha Lake | |
| Namibia2 | xxxxxx | xxxxxx | xxxxxx | *Barbus paludinosus* | Namibia | Hardap dam | |
| SA_Limpop | xxxxxx | xxxxxx | xxxxxx | *Barbus anoplus* | South Africa | Nwanedi Dam | |
| Namibia1 | xxxxxx | xxxxxx | xxxxxx | *Barbus paludinosus* | Namibia | Hardap dam | |
| SA_Buff | xxxxxx | xxxxxx | xxxxxx | Barbus anoplus | South Africa | Buffelskloof Spruit river | |
| Bn159 | xxxxxx | xxxxxx | xxxxxx | Barbus anoplus | South Africa | Buffelskloof Spruit river | |
| NZ2 | xxxxxx | xxxxxx | xxxxxx | *Gobiomorphus breviceps* | New Zealand | | |
| NZS | xxxxxx | xxxxxx | xxxxxx | *Gobiomorphus breviceps* | New Zealand | | |
| RU3Ab | EU241212 | EU241252 | | *Abramis brama* | Russia | Rybinsk | |
